# A phenotypic screen using splitCas9 identifies essential genes required for actin regulation during host cell egress and invasion by *Toxoplasma gondii*

**DOI:** 10.1101/2021.09.24.461619

**Authors:** Wei Li, Janessa Grech, Johannes Felix Stortz, Matthew Gow, Javier Periz, Markus Meissner, Elena Jimenez-Ruiz

## Abstract

Apicomplexan parasites, such as *Toxoplasma gondii*, possess unique organelles, cytoskeletal structures, signalling cascades, replicate by internal budding within a specialised compartment and actively invade and exit the host cell, to name a few aspects of the unique biology that characterise this phylum. Due to their huge phylogenetic distance from well established model organisms, such as opisthokonts, comparative genomics has a limited capacity to infer gene functions and conserved proteins can fulfil different roles in apicomplexans. Indeed, approximately 30% of all genes are annotated as hypothetical and many had a crucial role during the asexual life cycle in genome-wide screens. While the current CRISPR/Cas9-based screens allow the identification of fitness conferring genes, only little information about the respective functions can be obtained. To overcome this limitation, and to group genes of interest into functional groups, we established a conditional Cas9-system in *T. gondii* that allows phenotypic screens. Using an indicator strain for F-actin dynamics and apicoplast segregation, we identified critical genes required for defined steps during the asexual life cycle. The detailed characterisation of two of these candidates revealed them to be critical for host cell egress and invasion and to act at different time points in the disassembly of the intravacuolar F-actin network. While the signalling linking factor (SLF) is an integral part of a signalling complex required for early induction of egress, a novel conoid protein (conoid gliding protein, CGP) acts late during egress and is required for the activation of gliding motility.

## Main

Apicomplexans are early branching eukaryotes related to ciliates and dinoflagellates, with unique adaptations to an intracellular, parasitic existence. The huge phylogenetic distance to well established model organisms is also reflected by the fact that many genes are unique and annotated as hypothetical. With the advancement of the CRISPR/Cas9 technology in *T. gondii*, genome-wide screens allowed the identification of genes that are important during the asexual stage of the parasite^1^ with many hypothetical genes being critical for the survival of the parasite. While pooled screens allow the identification of fitness conferring genes, downstream assays are required to define functional groups based on the specific phenotype caused by their deletion.

For the adaptation of phenotypic screens in *T. gondii*, we previously attempted to implement a conditional Cas9 system based on ddFKBP^2^ and, while it was possible to identify genes critically involved in nuclear mRNA export, the ddFKBP-system^3^ suffered from inefficient regulation, prohibiting tight temporal control of Cas9 activity^2^. This resulted in non-essential genes being rapidly lost in a transfected pool (unpublished).

Here, we adapted a tight control system for Cas9, based on splitCas9 (sCas9)^4^. We generated indicator parasites, expressing chromobodies directed against F-actin^5^ and a marker for the apicoplast (FNR-RFP; a plastid like organelle) that allowed the identification of specific phenotypes in an image-based screen for parasites with abnormal invasion, replication and host cell egress and in parallel the characterisation of changes in F-actin dynamics and apicoplast segregation. We screened a library of 320 genes, with more than 40% annotated as hypothetical and conserved within the phylum of Apicomplexa^1^. Parasite mutants were grouped into defined phenotypes, focusing on changes in F-actin dynamics and delays in host cell egress. We identified several genes critical for host-cell invasion, replication and egress. Among them, we identified a novel conoid associated protein (TGGT1_240380) and a putative neurotransmitter symporter (TGGT1_208420) that are critical for distinct, independent steps during host cell egress and act at different times. Since, both proteins are highly conserved within apicomplexan parasites, they likely fulfil conserved, critical functions during host cell egress and invasion.

## Results

### Adaptation of a conditional Cas9-system in T. gondii

In order to achieve tight, temporal control of Cas9, we chose the sCas9 system^4^, where the Cas9 enzyme is split into two subunits (N- and C-terminus), which are fused to a FKBP and FRB domain. Upon addition of rapamycin these subunits interact and Cas9 activity is restored (Fig.1a). We generated transgenic *RHΔHX* parasites expressing both sCas9 subunits (Supplementary Fig.1a,b), resulting in the recipient strain RHsCas9.

**Figure 1.**
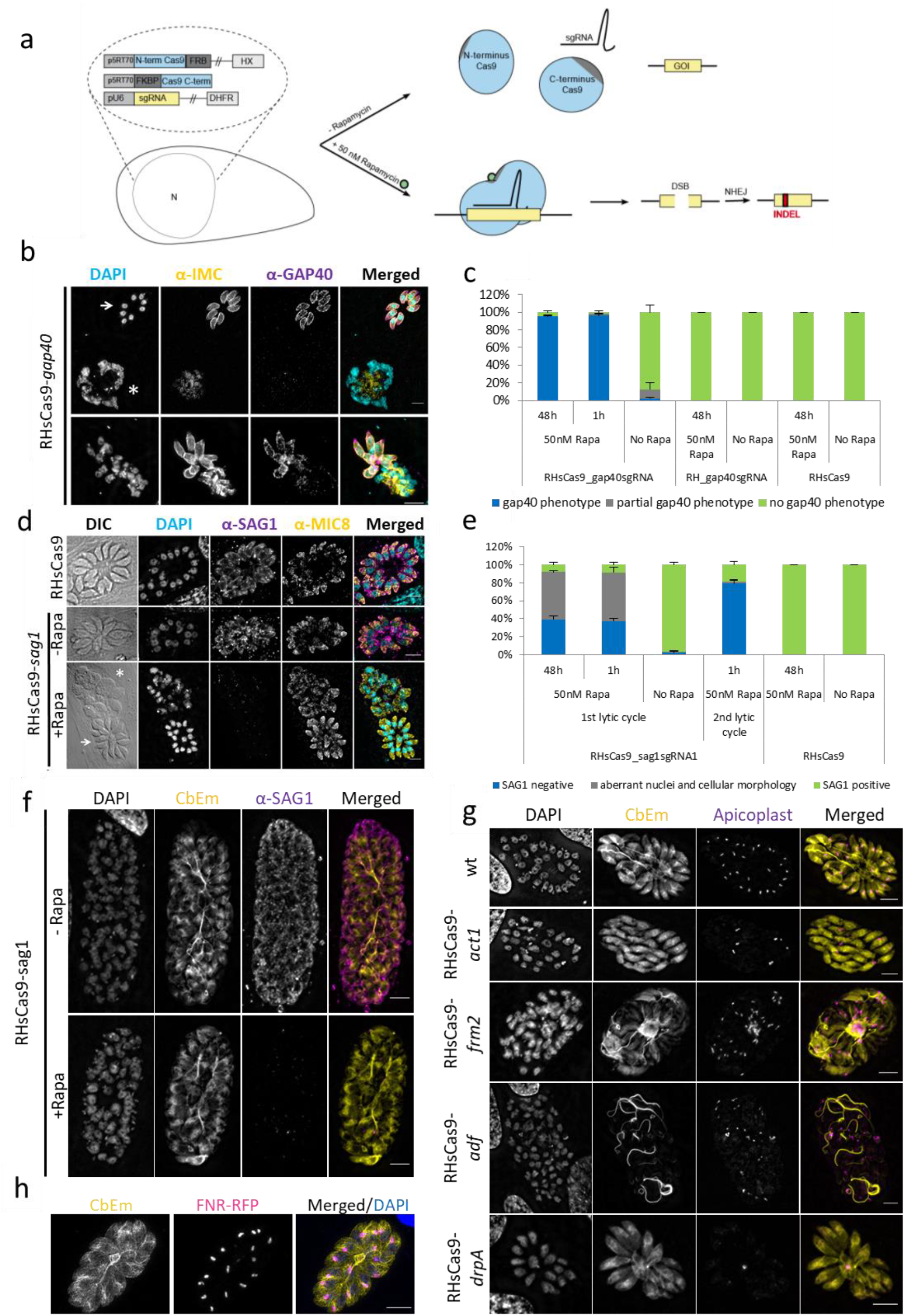
Adaptation and characterisation of the splitCas9-system in *T. gondii.* **a,** Schematic of the splitCas9-system. Transgenic parasites are generated co-expressing the two splitCas9 subunits together with a single-guide RNA (sgRNA). Upon addition of rapamycin the two subunits dimerise, leading to reconstituted Cas9 activity and therefore to disruption of the gene targeted by the sgRNA. **b,** Analysis of RHsCas9-gap40 parasites that were treated with 50 nM rapamycin for 48h before fixation using indicated antibodies. Scale bars are 5μm. Three distinct phenotypes can be observed: the gap40 phenotype, as described previously^7^ with collapsed IMC and loss of GAP40 expression (top panel, asterisk); a phenotype where some parasites within the PV are aberrant (bottom panel); and parasites with normal IMC and GAP40 localisation (top panel, arrow). **c,** Quantification of gap40 phenotypes shown in (a) in indicated parasite strains. Parasites were induced with or without rapamycin for the indicated time and fixed 48 h post infection. Only parasites expressing both the gap40sgRNA and the sCas9 system presented a gap40 phenotype after induction. Parasites were treated with rapamycin for 1h or the whole growth period of 48h as indicated. Data represents three independent experiments. For each condition 100 vacuoles were counted (total n=300). Average and standard deviation (SD) are represented. **d,** Analysis of RHsCas9-sag1 parasites that were treated with 50 nM rapamycin for 48h before fixation using indicated antibodies. Nuclei were stained with DAPI. Scale bars are 5μm. Three distinct phenotypes can be observed: healthy vacuoles lacking SAG1 (bottom panel, arrow); and parasites lacking SAG1 while displaying aberrant nuclear and cellular morphology (bottom panel, asterisk). **e,** Quantification of the phenotypes shown in (d). Aberrant nuclei and cellular morphology were observed only when *sag1* was disrupted (KO) by sCas9 activation (1^st^ lytic cycle). Abundance of non-healthy parasites was reduced to background levels when induced RHsCas9-*sag1* parasites were mechanically lysed, transferred onto fresh host cells and grown again for 48h in a second lytic cycle (2^nd^ generation; total incubation of 96h). Parasites were treated with rapamycin for 1h or for the whole growth period of 48h as indicated. Data represent three independent experiments. For each condition, 100 vacuoles were counted (total n=300). Average and standard deviation (SD) are represented. **f,** Analysis of indicator parasites expressing sgRNA targeting *sag1.* Disruption of *sag1* has no effect on the F-actin network. Scale bars are 5 μm. **g,** Analysis of indicator parasites expressing indicated sgRNAs. IFA depicting the effect of *drpA* (RHsCas9-*drpA*), or *adf* (RHsCas9-*adf*), *act1* (RHsCas9-*act1)* and *frm2* (RHsCas9-*frm2*) disruption on actin network and apicoplast segregation. To achieve gene disruption (KO), parasites were incubated with 50 nM rapamycin for 1h and fixed after 48h.Nuclei were stained with DAPI. Scale bars are 5 μm. **h,** Depiction of indicator parasites co-expressing CbEm, FNR-RFP and the sCas9-subunits. Scale bar is 5 μm.

To test efficiency and specificity of this system, we generated vectors for stable expression of small guide RNAs (sgRNA) against previously described essential and non-essential genes, such as *sag1*^6^ and *gap40*^7^. These vectors were randomly integrated into the genome of RHsCas9 and into *RHΔHX* parasites (Supplementary Table1; Supplementary Fig.1c,d,e). Subsequently, parasites were induced with 50nM rapamycin for 1h or 48h before they were analysed. Upon rapamycin treatment, a *gap40-like* phenotype in RHsCas9-gap40 parasites, as described in the literature^7^, was observed in up to 95% of vacuoles, but not in RH-gap40 or RHsCas9 strains (Fig.1b,c). No difference in induction rate was observed upon 1 or 48 hours of rapamycin treatment, demonstrating that the sCas9 system allows efficient and rapid activation (Fig.1c).

A second control experiment targeted the non-essential surface antigen 1 (SAG1)^6^. While >95% of parasites demonstrated loss of SAG1, we found that up to 50% of parasites also showed aberrant morphology of their nuclei in addition to loss of SAG1 (Fig.1d,e). This phenotype was only present in rapamycin-treated parasites co-expressing sCas9 and *sag1* sgRNA. No impact on parasite morphology was observed in RHsCas9 parasites (cultured with or without rapamycin), indicating that this phenotype is not caused by general toxicity of sCas9 (Fig.1e). We reasoned that the introduction of a double strand break (DSB) in the genome leads to a replication defect in a subpopulation of induced parasites.

To assess this phenotype further, we induced RHsCas9-sag1 parasites for 1h, mechanically released them from the host cell 48h later and allowed them to infect fresh HFF cells, thus representing the 2^nd^ lytic cycle. Most of these parasites were negative for SAG1 (79% ±4.8), and no aberrant morphology or nuclei were detectable (Fig.1e). To address if the nuclear phenotype is caused by the introduction of a DSB, we isolated RHsCas9-*sag1*KO and confirmed specific gene disruption (Supplementary Fig.2a). Next, we silently mutated *sag1* into a non-cleavable copy of *sag1* (*sag1\**) and introduced it into RHsCas9-*sag1*KO and non-induced RHsCas9-sag1 parasites. (Supplementary Fig.2b). Upon induction with rapamycin, we found no reduction of SAG1-positive parasites could be observed, since both strains expressed the non-cleavable *sag1*\*. However, the parasite strain with intact endogenous *sag1* (RHsCas9-sag1) demonstrated a similar aberrant phenotype as described above, while no phenotype could be observed in case of RHsCas9-*sag1*KO containing only the non-cleavable *sag1\** (Supplementary Fig.2c,d). This demonstrates that the introduction of a DSB can cause an aberrant nuclear phenotype.

In summary, the sCas9 system allows the efficient generation of conditional mutants, but one needs to ensure that the phenotype of interest is sufficiently distinct from the aberrant nuclear (non-specific) phenotype (Fig.1d,e) that needs to be subtracted in the readout of any screen.

### Generation of an indicator parasite line for a forward genetic screen for F-actin and egress mutants

To assess if specific phenotypes related to changes in F-actin dynamics and apicoplast replication can be identified, we tested a sCas9 strain expressing the actin Chromobody CB-EmeraldFP (CbEm)^5^ by disrupting *sag1*, as negative control, *drpA*^8^, which should only affect apicoplast division, *act1*^5,9^, *adf*^10^ and *formin 2 (frm2)*^11^ (Fig.1f,g and Supplementary Fig.1g), which should cause different effects on F-actin dynamics and apicoplast division. For the phenotypic screen, we included the apicoplast marker, FNR-RFP^12^ (Fig.1h), to facilitate image acquisition.

Stable transfection with the respective sgRNA expression vectors and disruption of the target gene was validated by sequencing of the respective loci after gene disruption (Supplementary Fig.1g). Parasitophorous vacuoles (PV) containing parasites with abnormal nuclei, representing the non-specific phenotype shown above, were excluded from subsequent analysis. As expected, disruption of *sag1* or *drpA* had no detectable effect on the formation of the intravacuolar F-actin network (Fig.1g) and a strong defect in apicoplast segregation was observed in the case of *drpA*-disruption, as previously described^8^. In contrast, disruption of *act1* and *adf* led to disintegration and stabilisation of the intravacuolar network respectively (Fig.1g) as described^5, 9, 13^. In addition, disruption of the actin nucleator FRM2 also led to stabilisation of the intravacuolar network and abrogation of the intracellular polymerisation centre, replicating the effect observed when excising the locus using a DiCre strategy^11^. In all cases, apicoplast inheritance was affected as expected. Therefore, the indicator strain allows the identification of specific phenotypes to apicoplast maintenance, actin dynamics, or both. Finally, we assessed if delays in parasite egress can be identified in order to screen for genes required for host cell egress (Supplementary Fig.3). In good agreement with the literature, disruption of *adf* and *act1* led to a significant delay in egress, while the behaviour of parasites with disrupted *sag1* was similar to WT parasites (Supplementary Fig.3a,b,c).

### Phenotypic screening using a custom-designed sgRNA library

For the phenotypic screen, we curated a library containing hypothetical genes without a signal peptide that are conserved in apicomplexans and have been hypothesised to be fitness conferring (phenotypic score <-1.5)^1^. In total 320 genes were targeted (Supplementary Table2). As internal controls sgRNAs targeting *gap40* (parasite replication^7^) and *prf* (F-actin dynamics and egress^14^) were included (Supplementary Fig.4a,b).

Synthesised sgRNAs were cloned into a vector containing the DHFR selection cassette as previously described^15^. The number of bacteria in the recovered sample carrying ligated plasmid was estimated to be 3 × 10^6^ cfu, exceeding the number recommended to maintain complexity^15^. Library complexity was confirmed by sequencing sgRNA from 35 colonies picked at random (not shown), out of which we identified 29 unique gRNAs (83%).

Next, the library was transfected into the indicator strain, followed by drug selection, and sorting of single parasites into ten 96-well plates (Fig.2a).

**Figure 2.**
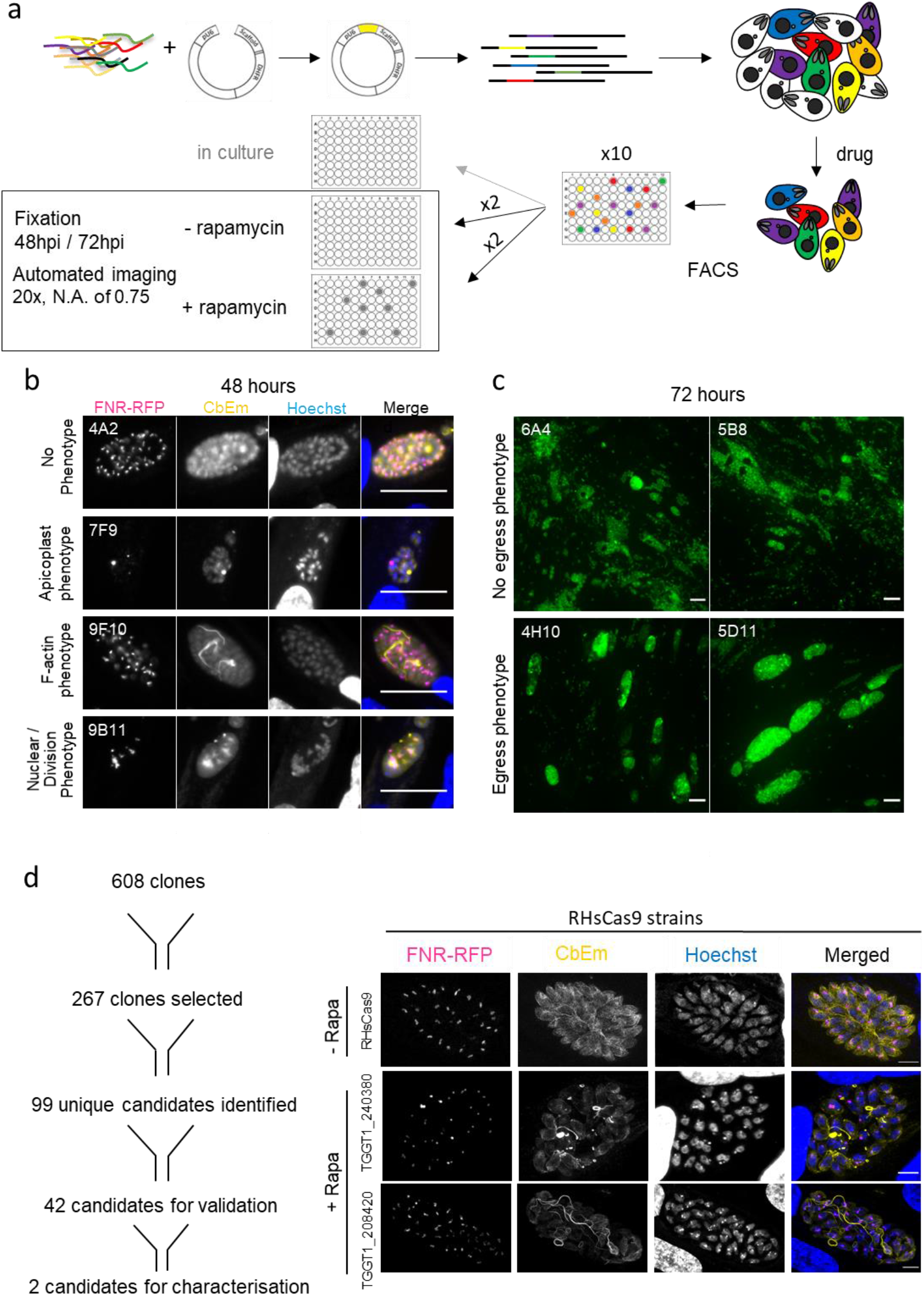
Phenotypic screen for actin dynamics, apicoplast segregation and egress mutants. **a,** Scheme of the experimental design. Indicator parasites, RHsCas9-CbEm-FNR-RFP, were transfected with the sgRNA library and grown in human foreskin fibroblasts (HFFs). Parasites were selected with pyrimethamine and sorted into 96-well plates. After 7 days, they were split into replica plates. Two replica plates were induced for 48h or 72h with rapamycin, before automated imaging was performed. Candidate clones were subsequently picked from non-induced replica plates. **b,** Representative images of observed phenotypes 48h post induction. Scale bars are 25 μm **c,** Representative images 72 hpi. Top row shows clones without egress phenotype. Bottom row shows two examples of clones with a strong egress phenotype. Scale bars, 30 μm **d,** Schematic representation of the selection of candidate genes (left) and phenotype of the two identified egress mutants 48 hpi. Note that the apicoplast appears normal, while slight differences in the F-actin network, which appears more prominent and condensed, can be observed. Parasites were induced for 48 h with rapamycin before fixing and imaging. Scale bars, 5 μm.

We obtained 608 clonal parasites that were inoculated onto three replica plates. Two sets were induced with rapamycin for 48 and 72h, followed by fixation and automated imaging. Images were independently analysed twice by different authors and graded by the relative strength of phenotypes (Fig.2b,c). From this analysis, a total of 267 clones, were put forward for analysis of integrated sgRNA. We dismissed clones with integration of multiple different sgRNAs, which caused “super-aberrant” phenotypes (Supplementary Table3, Supplementary Fig.4c). Importantly, our internal controls were isolated multiple times (Supplementary Table3, Supplementary Fig.4a,b), demonstrating the robustness of this screening approach. This procedure resulted in a total of 99 candidate genes (Supplementary Table3) that were prioritised by re-analysing the obtained images. We focused on clones with the highest induction rates and excluded candidates with strong replication defect (see above; Fig.2b,c). In summary, disruption of 42 genes showed detectable differences in F-actin formation, apicoplast segregation and/or host cell egress (Supplementary Table4). However, it should be noted that F-actin changes were relatively minor, when compared to the internal control (profilin) (Supplementary Fig.4a) or the effects seen upon disruption of *adf or frm2* (Fig.1h). We succeeded in tagging 12 of the 16 candidates classified as F-actin and/or apicoplast phenotype (Supplementary Fig.5). Interestingly, one of these genes, TGGT1_208420, localised to the intravacuolar network and the apical pole of the parasite and was also found as important for natural egress (see below; Fig.3a and Supplementary Fig.7a).

**Figure 3.**
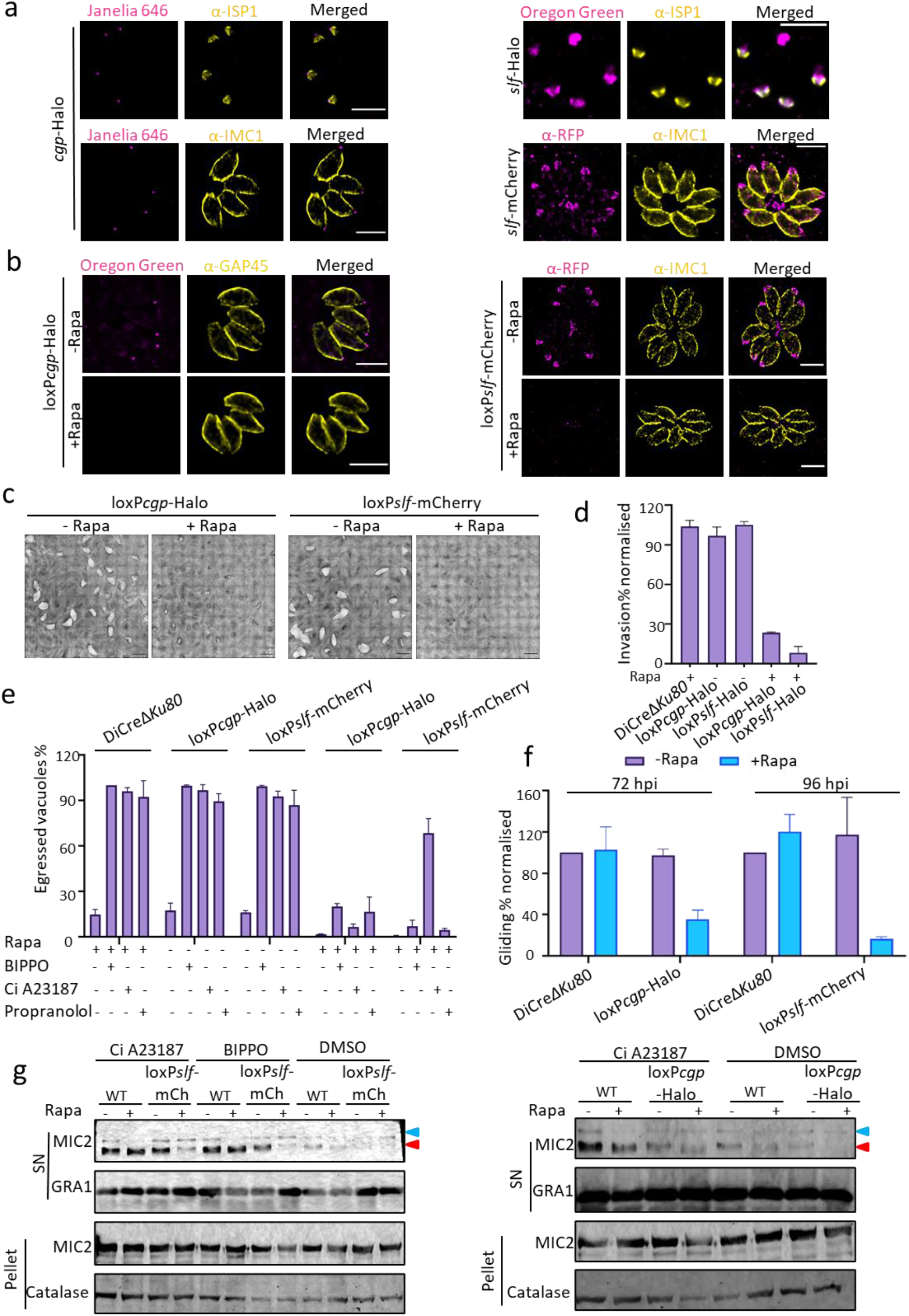
Analysis of *cgp* and *slf.* **a,** Endogenously-tagged TGGT1_240380 (*cgp*-Halo) localised at the apical tip, while TGGT1_208420 (*slf*-Halo or *slf*-mCherry) demonstrated a dual localisation at the apical region and the residual body. Dual labelling was performed to stain the apical cap (ISP1) of the parasite. Scale bar: 5 μm. **b,** Analysis of conditional knockouts for *cgp* and *slf* using the DiCre-system. IFA depicts floxed *cgp*-Halo (loxP*cgp*-Halo) and floxed *slf*-mCherry (loxP*slf*-mCherry) induced or not with 50 nM rapamycin (Rapa). 72 hours post induction, both proteins were not detectable by IFA. Dual labelling was performed to stain the IMC (GAP45 or IMC1) of the parasite, which shows no differences upon deletion of *slf* or *cgp.* Scale bar, 5 μm. **c,** Plaque assays of loxP*cgp*-Halo and loxP*slf*-mCherry parasites confirm a severe growth defect upon deletion of *slf* or *cgp*. Parasites were treated ± 50 nM rapamycin for 6 days before fixation. Scale bar,1.5 mm. **d,** Invasion-attachment assay for the indicated parasites lines. Results were normalised to DiCreΔ*ku80* strain. This assay was performed in triplicate. Bars represent mean ± standard deviation (SD). For each condition, at least 150 vacuoles were counted (total n≥450). **e,** Induced egress assay in the presence or absence of different inducers: Calcium ionophore (Ci) A23187 (2 μM) for 5 min, BIPPO (50 μM) for 5 min, and Propranolol (125 μM) for 7 min. Experiment was performed in triplicates. Data shown as Mean ± SD. For each condition 100 vacuoles were counted (total n=300). **f,** Quantification of trail deposition. Results were normalised to DiCreΔ*ku80* strain. 3 biological replicates were performed. Data were presented as mean ± SD. **g,** Microneme secretion assay was perform on wildtype (WT) parasites, loxP*cgp*-Halo and loxP*slf*-mCherry (loxP*slf*-mCh). While parasites lacking CGP showed normal microneme secretion comparable to WT, *slf* cKO parasites had a decreased secretion. Triangles indicate the unprocessed (blue) and processed (red) form of MIC2. This assay was performed in 3 biological replicates. Representative images are shown.

### Selection of candidates potentially involved in parasite egress

Next, parasite egress was analysed 72h after inoculation and induction of sCas9. At this time point, most initially infected host cells were lysed and parasites reinvaded neighbouring host cells (Fig.2c). We identified 33 clones with a potential delay in host cell egress (Supplementary Table 4). Upon a second round of visual inspection, parasites forming smaller or aberrant PVs, indicating obvious replication defects, were excluded, resulting in the identification of candidates where conditional disruption resulted in a delayed egress phenotype (Supplementary Table4).

Analysis of stimulated egress using calcium ionophore A23187 (Ci A23187) revealed that disruption of 4 candidate genes resulted in significantly delayed egress. In contrast, control parasites with disrupted *act1* and *adf*, were inhibited in egress (Supplementary Fig.3a and Supplementary Fig.6a,b). Since disruption of the gene TGGT1_252465, resulted in slower replication, it was excluded from additional analysis (Supplementary Fig.6c). Finally, the 3 remaining candidates also demonstrated a significant reduction in host cell invasion rates when compared to controls (Supplementary Fig.6d). One candidate (TGGT1_248640) has recently been described as one of the non-discharge proteins, TgNd6^16^.

Here we focus our analysis on the remaining 2 candidates, TGGT1_240380 and TGGT1_208420, named conoid gliding protein (CGP) and signalling linker factor (SLF) respectively (Fig.2d).

### CGP and SLF are required for egress and invasion

We tagged both genes C-terminally with Halo-Tag (*cgp*-Halo) or mCherry (*slf*-mCherry or *slf*-Halo) (Fig.3a, Supplementary Fig.7a). Both proteins showed an apical localisation, as evidenced via colocalisation with the apical IMC-marker ISP1^17^. In addition, SLF also accumulated in the intravacuolar network that connects individual parasites within the PV^5^ (Supplementary Fig.9a).

To validate the phenotype seen with sCas9, genes were floxed in the RHDiCreΔ*Ku80* strain^18^ (Fig.3b; Supplementary Fig.7, Fig.8a and Fig.9b) to generate conditional null mutants (cKO). Induction of DiCre using rapamycin led to a severe growth defect, as evidenced by plaque assays, confirming their critical role during the asexual life cycle of the parasite (Fig.3c). Further analysis fully confirmed the phenotype obtained using sCas9 and demonstrated a crucial function of both proteins in host cell egress and invasion, while parasite replication, morphology, apicoplast and secretory organelles were not affected (Fig.3d,e; Supplementary Fig.8b,d,e,f and Fig.9c,d,g,h,i).

Gliding motility was significantly reduced in both mutants (Fig.3f; Supplementary Fig.8c, Fig.9e; Movie_S1). Interestingly, in the case of parasites lacking *slf* (*slf*-cKO) gliding motility and egress (Fig.3e; Supplementary Fig.9e,f) could be partially rescued upon addition of Ci A23187 (Supplementary Fig.9f; Movie_S2), indicating that it is involved in a signalling cascade, upstream of intracellular calcium release. Indeed, *slf*-cKO showed a significantly reduced secretion of micronemes (Fig.3g), that can be partially rescued upon addition of Ci A23187, but not BIPPO^19^. Therefore, the phenotype of *slf*-cKO appeared similar to the phenotypes observed upon deletion of components of the phosphatidic acid (PA) signalling platform, such as diacylglycerol kinase 2 (DGK2), cell division control 50 related protein (CDC50.1), guanylate cyclase (GC) or unique GC organiser (UGO)^20^.

In the case of *cgp* knockout parasites (*cgp*-cKO), egress was blocked irrespective of addition of Ci A23187, BIPPO or Propanolol (Fig.3e). Furthermore, no defects of microneme secretion could be observed (Fig.3g), placing this protein into a different functional category.

### Analysis of host cell egress reveals that CGP and SLF act at two distinct, temporally controlled steps

We then characterised the sequential action of CGP and SLF during host cell egress. The intravacuolar network is rapidly disintegrated early in the egress process^5^, thereby acting as an early indicator for initiation of host cell egress. Therefore, we introduced CbEm^5^ into the UPRT locus of both mutants to analyse localisation and dynamics of F-actin (Fig.4a; Supplementary Fig.7k,l; Movies_S3, S4 and S5).

**Figure 4.**
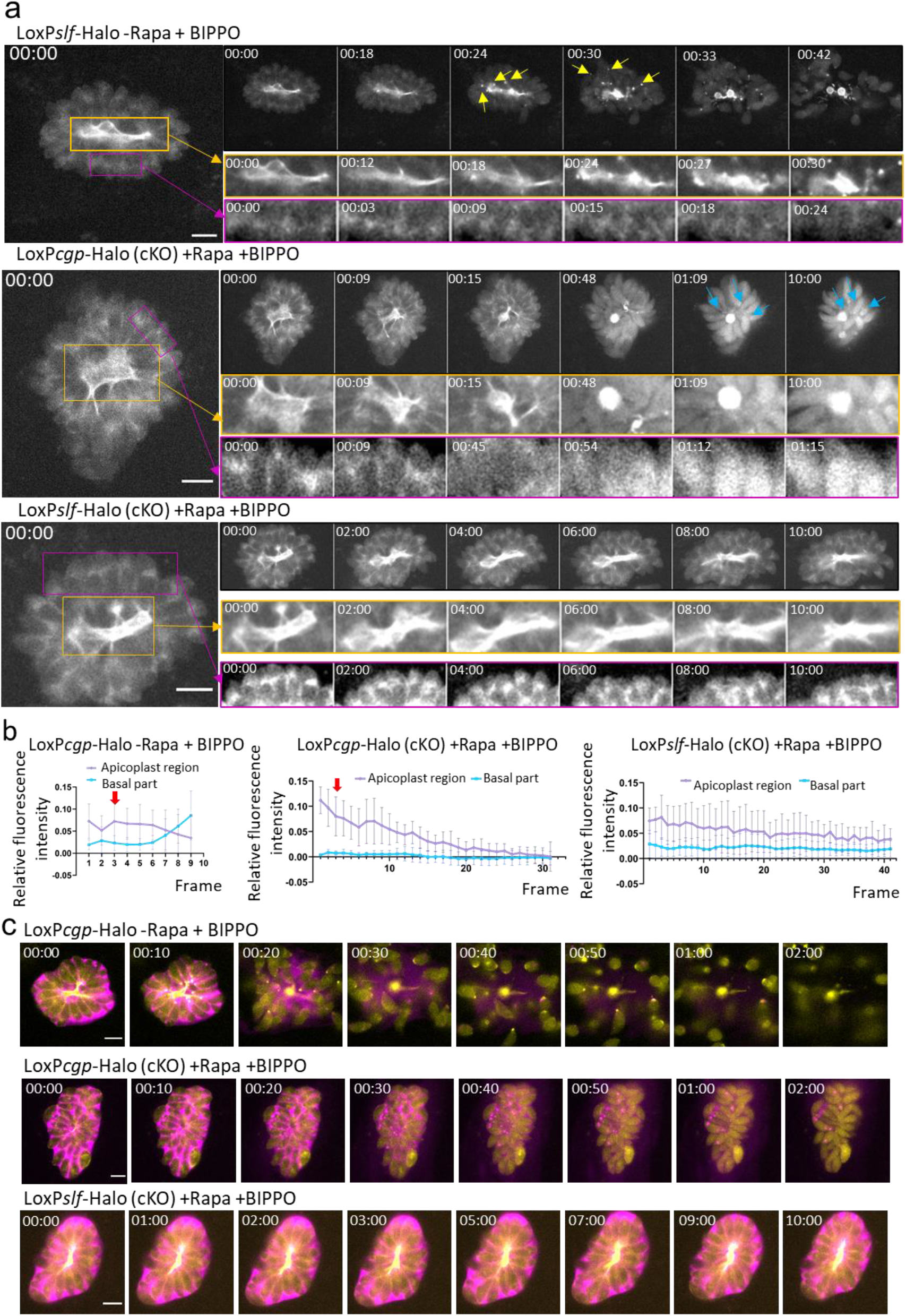
SLF and CGP act at different times during egress. **a,** Egress of parasites expressing CbEm labelling F-actin was induced with 50 μM BIPPO and imaged with an interval of 3 seconds between frames. yellow boxes show F-actin in intravacuolar network. Pink boxes show F-actin polymerisation centre close to the apicoplast/Golgi region (insets have enhanced contrast for better visualisation). Upper panel showing non-induced parasites as control (see movie_S4). F-actin disintegrated rapidly upon addition of BIPPO accompanied with F-actin accumulation at the basal pole (indicated by yellow arrows). The polymerisation centre close to the apicoplast/Golgi region appeared to become inactive during egress. Middle panel depicting F-actin in loxP*cgp*-Halo parasites previously induced with 50 nM rapamycin. Only parasites lacking signal for CGP were recorded (see movie_S3). Network disintegration and inactivation of the F-actin polymerisation centre close to the apicoplast/Golgi appeared normal, while parasites remained immotile and no relocalisation of F-actin to the basal pole is evident (blue arrow). Bottom panel depicting F-actin in loxP*slf*-Halo parasites previously induced with 50 nM rapamycin. Only parasites lacking signal for SLF were recorded (see movie_S4). The F-actin network remained intact and the F-actin polymerisation centre close to the apicoplast/Golgi remained active. Time is displayed in minutes : seconds. Scale bar, 5 μm. **b,** Quantification of the average relative fluorescence intensity of indicated parasites after induction of egress with 50 μM BIPPO. The graphs show the average of indicated individual measurements. Red arrows indicate the time where the F-actin network started to depolymerise. In control parasites disassembly of the F-actin network was followed by a reduction of polymerisation at the centre close to the apicoplast/Golgi region and an increase accumulation of F-actin at the basal pole of the parasite. Deletion of *cgp* resulted in similar behaviour of the F-actin network and comparable reduction at the polymerisation centre close to the apicoplast/Golgi, but no accumulation of F-actin at the basal pole. Parasites lacking *slf* showed no reorganisation of Factin upon stimulation of egress with BIPPO. Time interval between each frame is 3 seconds. **c,** Egress of parasites expressing CbEm (yellow) and SAG1ΔGPI-dsRed (pink) was induced with 50 μM BIPPO. Upper panel showing non-induced parasites, upon induction, dsRed signal diffused into the host cell, indicating lysis of parasitophorous vacuole membrane (PVM). Middle panel depicting parasites lacking CGP signal after rapamycin induction. Although the PVM lysed, parasites did not move out of the cell. Bottom panel depicting parasites lacking SLF signal after rapamycin induction, where dsRed signal is kept within the PV suggesting intact PVM. Time is displayed in minutes : seconds. Scale bar, 5 μm. See also movies_S5.

When analysing F-actin dynamics upon induction of egress, the following steps can be differentiated in WT parasites: 1) disassembly of the intravacuolar network, 2) reduction of F-actin nucleation close to the Golgi, probably caused by FRM2^5^) activation of motility and strong posterior accumulation of F-actin (Fig.4a,b). In the case of *slf*-cKO parasites, none of these steps could be observed indicating that initiation of egress is completely blocked and that the initial signals leading to the induction of microneme secretion are identical with regulation of F-actin dynamics during host cell egress (Fig.4a,c; Movie_S5). This phenotype was partially rescued upon addition of Ci A23187. (Movie_S4).

Interestingly, induction with propranolol led to a different egress phenotype, when compared to induction with BIPPO. Here the parasites that remained within the PVM were able to disassemble the filaments but appeared unable to initiate motility (Fig.4a; Movie_S4).

In contrast, deletion of *cgp* led to a block in a later stage during egress. While disassembly of the intravacuolar network and reduction of F-actin nucleation close to the Golgi occurred normally, neither posterior accumulation of F-actin nor parasite motility appears to be initiated (Fig.4a,b; Movie_S3).

In summary, this analysis highlights that SLF and CGP act at two temporally different steps during host cell egress. While SLF acts at the initiation step, CGP acts downstream, after the intravacuolar F-actin network has been disassembled.

Finally, we were interested to know if the PV-membrane (PVM) is dissolved. Therefore, we expressed *sag1*ΔGPI-dsRed, which is secreted into the PV. Upon lysis of the PV this protein diffuses into the cytosol of the host cell, as seen in case of control parasites (Fig.4c, top row). Using this assay, deletion of CGP had no influence on lysis of the PV, since dsRed signal diffused at a similar time as seen in case of controls. In contrast, upon deletion of SLF, the PVM remained intact and dsRed signal is trapped within the PV (Fig.4c; Movie_S5).

### SLF is required for integrity of the PA signalling complex

Bioinformatic predictions of SLF place this protein into the family of sodium neurotransmitter symporters and demonstrates that it is highly conserved in all apicomplexan parasites. This protein was previously identified as a putative and dispensable interaction partner of the signalling platform, since knockdown using the auxin-inducible degron (AID) system had no effect on the parasite lytic cycle^20^. However, disruption and excision of *slf* demonstrated one of the strongest phenotypes obtained in this screen, suggesting that the AID system is insufficient to knockdown protein levels of SLF. Indeed, SLF colocalised with other members of the PA signalling pathway such as GC, CDC50.1 and UGO (Fig.5a) at the apical tip of the parasite and the intravacuolar network. Importantly, deletion of SLF results in mislocalisation of other components of this signalling complex in the ER and vice versa (Fig.5b,c), indicating that this complex is assembled early in the secretory pathway, probably the ER, and only reaches its final destination if all partners are present. This was seen in 100% of vacuoles, where one component is missing. While this confirms an important structural role of SLF for functional assembly of the signalling complex, future studies are required to determine if this protein also acts as a sodium neurotransmitter as predicted in ToxoDB. In a first attempt, we focused on GABA (γ-aminobutyric acid) as a potential substrate for this putative symporter, since it was demonstrated that *T. gondii* synthesises high levels of GABA^21^ and modulates host cell migration using GABA as messenger^22^. However, we were unable to either complement the phenotype by adding increasing concentrations of GABA or to mimic the phenotype by addition of GABA analogues (Supplementary Fig.10). In conclusion, SLF is a critical component for the integrity of the PA signalling platform required for egress. Future analysis will reveal if this protein is directly involved in the signalling cascade by acting as a symporter.

**Figure 5.**
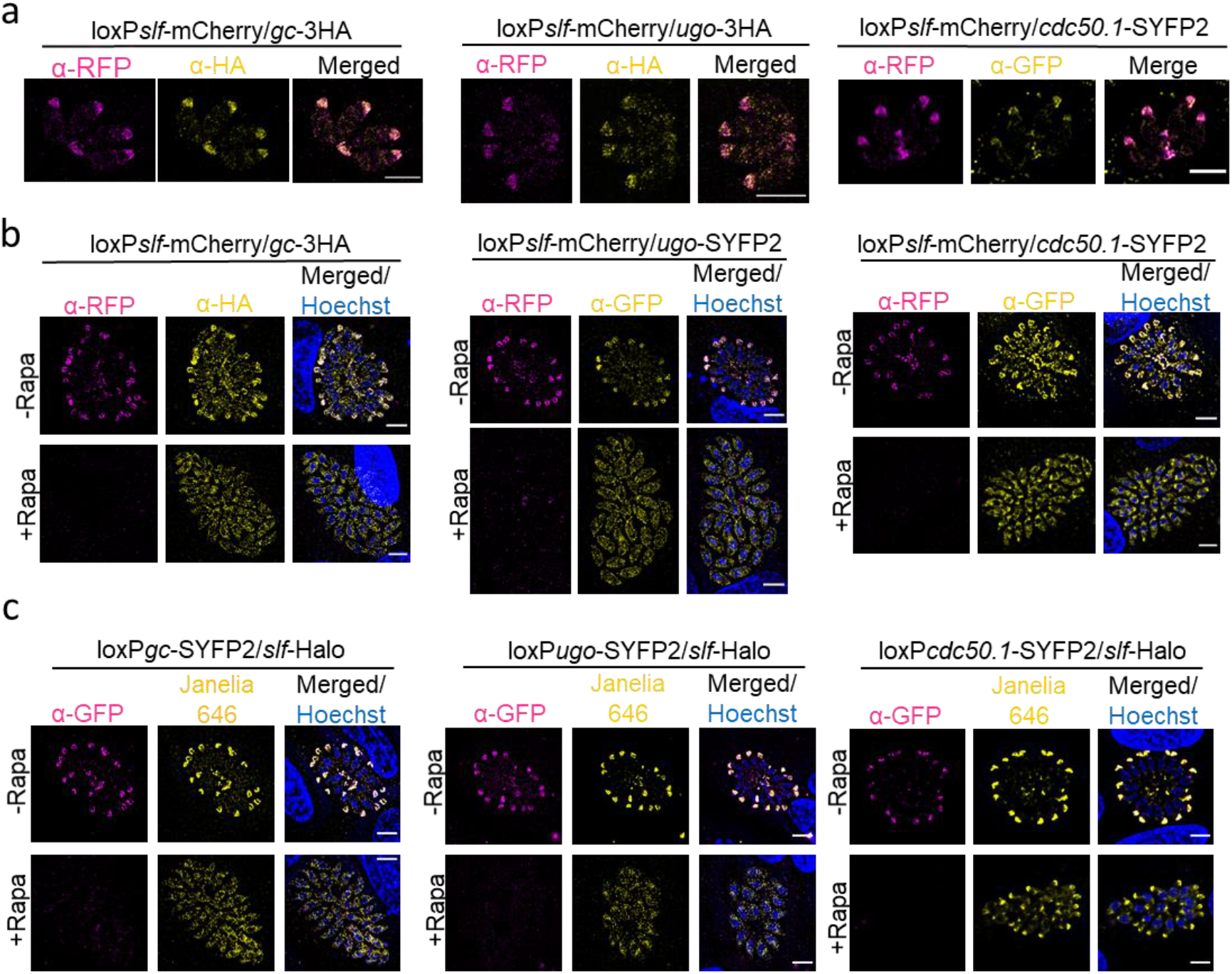
SLF is a crucial part of the signalling platform. **a,** GC, UGO and CDC50.1 colocalise with SLF. Scale bar: 5 μm. **b,** Localisation of indicated components of the signalling platform upon deletion of *slf.* Deletion of *slf-mCherry* results in mislocalisation of GC/UGO/CDC50.1. Analysis was performed 96 hours post induction. Scale bar, 5 μm. **c,** Deletion of *gc/ugo/cdc50.1* (see also Supplementary Fig. 7 for generation of conditional KOs for individual components) results in mislocalisation of SLF at 96 hours post induction. Scale bar: 5 μm.

### CGP is a novel component of the conoid

STED imaging showed a localisation of CGP anterior to the conoid markers RNG2, which localises to the second apical polar ring, and SAS6-like (SAS6L), a marker of the conoid body^23^. This apical localisation was detected in retracted and protruded conoids (Fig.6a). Importantly, conoid structure appeared unaffected upon deletion of *cgp* (Fig.6b) indicating that this protein does not play a key role for the integrity of the conoid itself. Similarly, the secretory organelles (micronemes and rhoptries) were not affected by deletion of *cgp* and secretion of micronemes occurred normally (Fig.3g; Supplementary Fig.8e,f).

**Figure 6.**
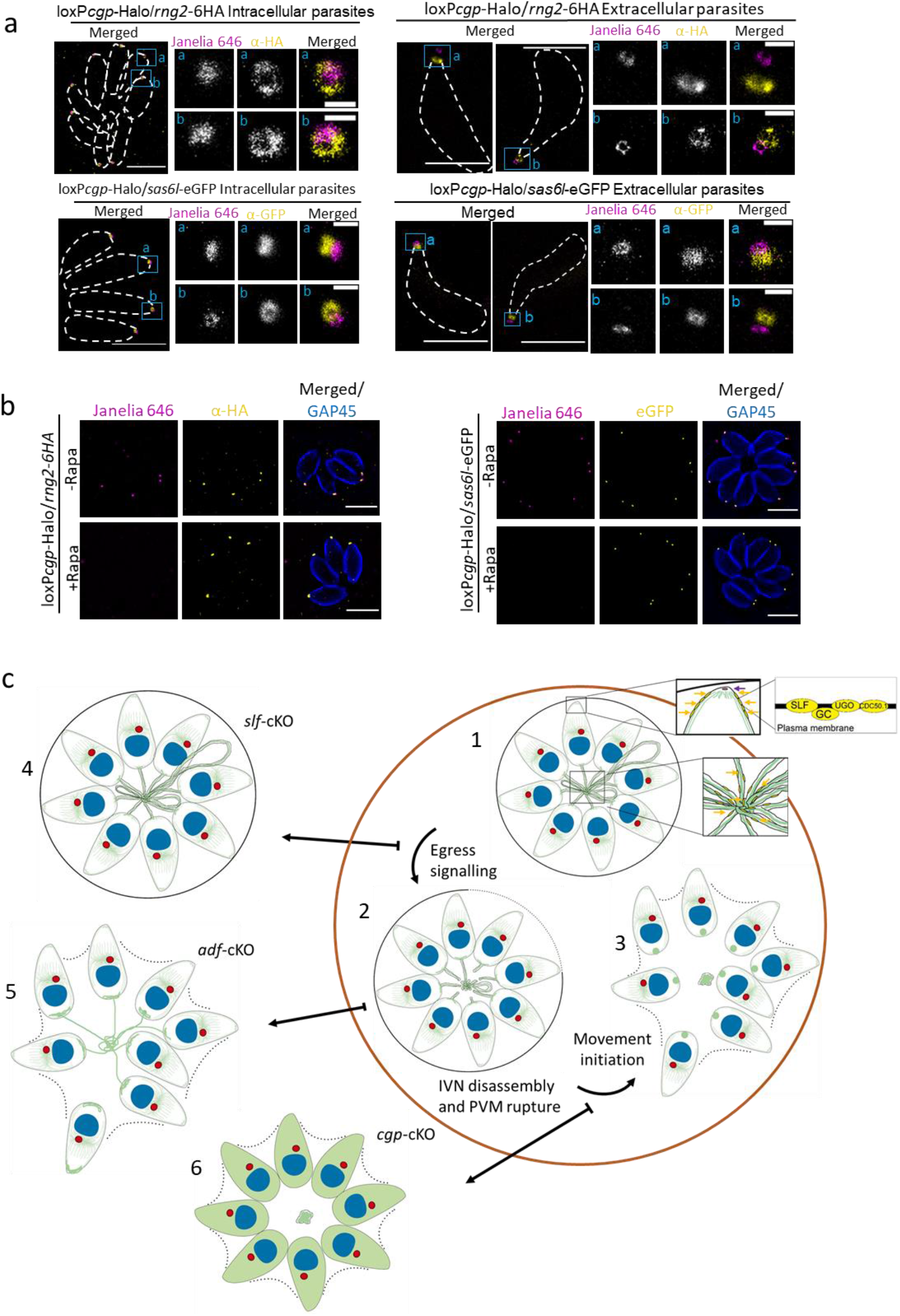
CGP localises to the preconoidal rings and egress models. **a,** STED microscopy of indicated parasites. Colocalisation of CGP-Halo with RNG2 or SAS6L proteins in intracellular parasites or extracellular parasites following conoid protrusion stimulation with 2 μM calcium ionophore A23187 for 10 min before fixation. White dash lines indicate parasite periphery. Scale bar, 5 μm for intracellular parasites and 3 μm for extracellular parasites. Scale bar of insets: 0.5 μm. **b,** Absence of CGP did not affect the localisation of RNG2 and SAS6L. Scale bar, 5 μm. **c,** Block of egress caused by deletion of SLF, CGP and ADF (see Supplementary Fig.3) is intrinsically linked to the disassembly of the F-actin network**. 1-3)** Natural egress process in a wildtype parasite. **1)** At the start of the egress the signalling platform consisting of CDC50.1, GC, UGO and SLF, localised at the plasma membrane of the apical tip and the intravacuolar network (IVN), initiates a cascade leading to egress. Zoom in boxes: yellow arrows indicate the position of the signalling platform at the apical tip and residual body. Purple arrow indicates the position of CGP at the conoid of the parasites. **2)** As a first step towards egress, the IVN disassembles, which coincides with lysis of the PVM. **3)** F-actin accumulates at the basal end of the parasite and motility is initiated. **4)** In the absence of SLF, the signalling platform is mislocalised and therefore not functional, resulting in early block of egress. No disassembly of the intravacuolar network or PVM lysis occurs. **5)** Depletion of actin regulatory proteins, such as ADF results in stabilisation of the network. Although parasites lyse the PVM and initiate motility, the network keeps connecting individual parasites resulting in delayed or blocked egress (see Figure S3 and ^11^). **6)** Deletion of *cgp* results in a late block of egress. The signalling cascade appears intact, leading to disassembly of the intravacuolar network and lysis of the PVM. Instead, no relocation of F-actin to the basal pole can occur and motility is not initiated.

While bioinformatic analysis of this protein suggests that it is highly conserved within apicomplexan parasites (not shown), no clear orthologue could be identified in other eukaryotes. Therefore, future studies are required to identify potential interaction partners and the mechanistic action during host cell egress.

## Discussion

In this study, we adapted an efficient and tightly regulated conditional Cas9-system based on sCas9^4^. Our detailed analysis of this system demonstrated that repair of DSB introduced by Cas9 is occasionally inefficient, leading to aberrant, non-specific phenotypes in a subpopulation of parasites. While these effects need to be taken into account when conducting a phenotypic screen, they can easily be deducted from the readout of the screen, especially when modern, automated image analysis methods are used that are now also available to determine the phenotype of *Toxoplasma-Host* interactions^24^. Screening of an indicator strain that co-expresses fluorescent actin binding chromobodies (CbEm)^5^ and an apicoplast marker^12^ allowed the identification of mutants with defects in apicoplast maintenance and F-actin dynamics in addition to mutants with inhibited host cell egress. Therefore, we curated a sgRNA library targeting 320 candidate genes that are conserved among apicomplexans. Based on the phenotypic characterisation of identified mutants, we did not identify a novel factor that is directly involved in regulation and organisation of F-actin dynamics during intracellular replication. Instead, several factors were identified, where F-actin dynamics was only slightly altered, when compared to positive controls (Profilin, ADF, Act1). In contrast, two novel genes identified here, *slf* and *cgp*, are crucially involved in the regulation actin dynamics during host cell egress where they play distinct roles.

Time-lapse microscopy analysis suggests that disassembly of the intravacuolar network precedes the initiation of motility and egress of parasites from the PV^5^ (Movie_S3, S4). Indeed, stabilisation of F-actin by depletion of ADF results in an egress phenotype, where parasites were able to initiate motility, but remain connected by the intravacuolar network (Fig.6c, Supplementary Fig.3).

When we applied this analysis to conditional mutants for SLF and CGP, we observed important differences in the behaviour during host cell egress. While deletion of SLF caused an early block in host cell egress, where neither the intravacuolar network nor the PVM is disassembled, deletion of CGP caused a late block in egress, where parasites disassembled both the intracellular F-actin network and PVM, but are incapable of initiating motility and leaving the host cell. While CGP is not involved in microneme secretion, it is likely that it directly or indirectly regulates gliding motility by regulating F-actin dynamics, since deletion of CGP resulted in clear differences in F-actin dynamics, as evidenced by missing posterior accumulation of F-actin.

In the case of SLF, we found that this protein is an important component of a previously described PA signalling platform required for host cell egress^20^, since deletion of SLF resulted in mislocalisation of the remaining components, CDC50.1, GC or UGO within the ER of the parasite. Together this indicates that the signalling complex needs to be assembled within the ER before being transported to the apical tip and intravacuolar network, where it fulfils its critical function.

These findings highlight the robustness of this approach for finding essential genes based on their function. In future screens, establishment of different indicator lines expressing markers for the secretory organelles, IMC or mitochondria could be used to specifically identify genes involved in organellar biogenesis, parasite replication or maintenance of the endosymbiotic organelles, to name a few examples for future applications of this technology.

Concurrently with our work, Smith and colleagues developed an alternative strategy, based on high-throughput CRISPR-mediated tagging of candidate genes with the AID-system and demonstrate the efficiency of their strategy by labelling and downregulation of the *T. gondii* kinome, resulting in the characterisation of kinases, involved in diverse functions. Both strategies are highly complementary and have a huge potential to screen the *T. gondii* genome in search for key candidates based on their function.

## Materials and Methods

### Cloning DNA constructs

The N- and C-termini of the Cas9 enzyme (split4 variant) were amplified from the original plasmids provided by Zetsche and colleagues^4^ via PCR. The PCR amplicons were ligated into the pGEM®-T Easy vector and sequenced. Subsequently, they were cloned into the *Toxoplasma* expression vector p5RT70-HX^26^ via the restriction enzymes EcoRI and PacI. For the C-term-Cas9 vector, the *hx* selection marker was removed by restriction with SacII. The resulting plasmids were confirmed by sequencing.

pU6-*sag1* gRNA-scaffold and *pU6-gap40* gRNA-scaffold sequence was synthesised and cloned into a backbone vector containing the DHFR resistance cassette by GeneScript. The Q5® Site-Directed Mutagenesis Kit (New England Biolabs) was used to insert sgRNAs targeting control genes of interest (*act1, adf, frm2* and *drpA*) into the synthesised sgRNA plasmid according to the manufacturer’s instructions. Importantly, a universal reverse primer was used together with a forward primer to which the whole sgRNA sequenced was attached, see also^27^. Further sgRNAs used in this study were cloned into the universal pU6 vector via BsaI digestion, primers annealing and standard ligation using T4 Ligase (see oligo sequences Supplementary Table 5) as previously described^27^. All sgRNA-plasmids were confirmed by sequencing.

To generate mutated *sag1\**, genomic DNA was amplified and inserted via EcoRI and PacI into p5RT70-HX^26^. Mutations at the sgRNA binding sequence were introduced via mutagenesis using Q5 Site Mutagenesis following manufacturer’s protocol (NEB) using primers described in Supplementary Table 5.

### Culturing of T. gondii and host cells

*T. gondii* tachyzoites were passaged onto Human foreskin fibroblasts (HFFs; ATCC, SCRC-1041) monolayers at 37 °C and 5% CO_2_ in DMEM (Sigma, D6546) supplemented with 10% FBS (BioSell FBS.US.0500), 4mM L-Glutamate (Sigma, G7513) and 20 μg/ml gentamicin (Sigma G1397).

### Generation of transgenic parasites

Freshly lysed *Toxoplasma* tachyzoites were transfected with Amaxa 4D-Nucleofector system (Lonza). ~1×10^6^ parasites were centrifuged and resuspended in 50 μl P3 buffer. Up to 20 μg of DNA for transfection, including vectors, donor DNA and/or single stranded DNA, was ethanol precipitated and resuspended in another 50 μl of P3 Buffer. Both resuspensions were mixed in a 100 μl cuvette (P3 Primary cells 4D-Nucleofector X kit L, V4XP-3024, Lonza). The programme FI-158 was used for electroporation. Immediately after transfection, parasites were resuspended in fresh complete DMEM and added onto a dish with confluent HFF cells.

For generation of sCas9-parasites, a total of 20 μg of the vectors containing the sCas9 subunits were linearised with NotI and transfected simultaneously in *RHΔhxgprt* (RHΔHX), adding NotI restriction enzyme to the transfection mix. Integrants were selected with 25 mg/ml mycophenolic acid (MPA) and 40 mg/ml of xanthine ^28^. After isolating a clone containing both functional subunits, sgRNAs against the *hxgprt* cassette were transfected and the parasites were selected with 80 mg/ml 6-Thioxanthine^28^. The *hxgprt* cassette was then sequenced to confirm the introduction of indels.

Vectors containing sgRNAs targeting control genes *(gap40, sag1, act1*, etc) were linearised via NotI and transfected into RHΔHXsCas9 as described above. To select for the *dhfr* resistance marker, parasites were treated with 1 μM pyrimethamine (Sigma 46706; Donald and Roos 1993). Insertion of sgRNAs were confirmed by PCR of the sgRNA cassette (Supplementary Fig.1f).

For the generation of RHsCas9-CbEm-FNR-RFP, RHsCas9 parasites were firstly transfected with the vector containing the CbEm cassette^5^. Transfectants were enriched via Fluorescence Assisted Cell Sorting (FACS; S3 BioRad), and a clone was isolated via limiting dilution. Secondly, the vector containing FNR-RFP was linearised via NotI and transfected into RHsCas9-CbEm strain and selected with 20 μg/ml chloramphenicol ^6^. After selection, clones were isolated by limiting dilution.

### sgRNA library generation and transfection

Guide RNAs were synthesised (CustomArray, Inc. USA), amplified by PCR, and cloned as a pool into a plasmid using Gibson Assembly (NEB E26115) as performed by Sidik et al. (2018)^15^. The vector plasmid carries a pU6 promoter, and a DHFR cassette. Assembled plasmids were transformed in two batches, and bacteria allowed to grow to log phase. The plasmids were extracted, purified, pooled, and 200 ng retransformed into the bacteria for further library amplification. Library complexity was estimated based on 1) the number of independent colonies obtained after transforming bacteria with 200 ng of library DNA (as described in Sidik et al. 2018^15^) and 2) random isolation of individual colonies and sequencing of the respective sgRNA. From the colony picking, 10 μg of different isolated gRNAs were transfected individually into RHsCas9-CbEm-FNR-RFP and analysed in parallel to the screen described below.

Following the generation of the vector library, 60 μg of pooled plasmid library was transfected into RHsCas9-CbEm-FNR-RFP line specifically created to be used as an indicator strain for this project. This transfection gave rise to a pool of parasites which were collectively carrying the guide RNA library. A minimum of 1×10^7^ parasites were passaged each time. To obtain clonal populations, transfectants were subjected to 3 weeks of pyrimethamine selection followed by FACS (BD FACSAriaIII) into ten 96-well plates, one event per well.

### Phenotypic screening of sCas9 mutants

Conditional mutagenesis via sCas9, was induced using 50 nM rapamycin for 48 and 72 h, and the plates were imaged using the LasX navigator on a Leica DMi8 widefield microscope attached to a Leica DFC9000 GTC camera, using a 20x objective. After choosing the correct carrier on the navigator, the plates were aligned and three random images were taken from each well, using the ‘On demand’ adaptive autofocus setting. The images obtained were independently screened by eye by two investigators before subsequent selection of the candidate clones. Clones which were seen to exhibit aberrant organellar morphologies or altered F-actin dynamics, and egress defects were then selected and the guide RNAs present in the clonal populations, which exhibited phenotypes deemed relevant to the project, were then sequenced to identify the gene disrupted (Supplementary Fig.4d). To prioritise the candidates for further characterisation, the phenotypes observed were graded from 1 (least severe) to 4 (most severe) (Supplementary table 3).

### Generation of tagged and floxed strains

Guide RNAs for cleavage upstream and downstream of the genes of interest were designed using EuPaGDT ^29^. Sequences of all sgRNAs employed in this study are detailed in Supplementary Table 1. The sgRNAs were ligated into a vector coding for Cas9-YFP expression, as has been previously described ^30^.

Repair templates for integration of a loxP sequence and a tag were generated as in Stortz et al. (2019)^11^. Briefly, repair templates carrying the upstream loxP sequence were ordered as ssDNA oligos (ThermoFischer Scientific), the loxP sequences being flanked by 33 bp of homology.

The repair templates carrying tags, such as 3xHA, SYFP2, Halo and SNAP, were generated by PCR where the 50 bp of homology flanking the tags were introduced via the primer. The repair templates were pooled according to the gene and purified using a PCR purification kit (Blirt) (Supplementary Fig.7a,b).

Parasite transfection, sorting and screening for positive mutants was done according to Stortz *et al.* (2019)^11^. Briefly, newly released RHDiCreΔ*ku80* tachyzoites^31^ were transfected with the repair templates and 10-12 μg of vectors (encoding Cas-YFP and the respective sgRNAs) as described above. The parasites were mechanically egressed 24 to 48 h after transfection, passed through a 3 μm filter, and those transiently expressing Cas9-YFP enriched via FACS (FACSARIA III, BD Biosciences) into 96-well plates (a minimum of 3 events per well). Resultant clonal lines were screened by PCR and repair template integration confirmed by sequencing (Eurofins Genomics).

For the insertion of CbEm into tagged lines, a specific sgRNA targeting the uracil phosphoribosyltransferase (UPRT) locus was designed and cloned into a Cas9-YFP-expressing vector. CbEm cassette from the original plasmid used in Periz *et al.*^5^, was PCR amplified and integrated into the UPRT locus (Supplementary Fig.7k).

### Immunofluorescence assays

Immunofluorescence analysis was carried out as previously described^32^. Briefly, parasites were fixed in 4% paraformaldehyde for 15-20 minutes at room temperature. Samples were blocked and permeabilised in phosphate-buffered saline (PBS) with 2% BSA and 0.2% Triton X–100 for at least 20 minutes. Antibody labelling was performed using the indicated combinations of primary antibodies for 1 h, followed by the incubation with secondary antibodies for another 45 min. All antibodies are listed in Supplementary Table 6. α-GFP-ATTO 488 (1:500, Nano Tag Biotechnologies, N0304-At488-L) was directly used for 1h after permeablisation. Parasites containing Halo or SNAP tags were incubated with specific dyes for 1 h and washed away, followed by incubation with media for 1 h before fixation unless specifically indicated elsewhere (see Supplementary Table 6). Images were taken using Leica DMi8 Widefield microscope or an Abberior 3D STED microscope. The library parasite pictures obtained using the Abberior 3D STED microscope were taken using the confocal setting for FNR-RFP and Hoechst, and STED for the CbEm imaging.

### Invasion/replication assays

24 h invasion/replication assays were performed using sCas9 parasite clones isolated as previously described, with some changes^32^. 5 × 10^6^ parasites were used to infect covers and were left to settle down on ice before allowed invasion for 20 min. Parasites were stained with α-SAG1 antibody without permeabilisation and α-GAP45 antibody after permeabilisation. For invasion, the number of vacuoles in 10 randomly selected fields of view were counted for each parasite line and condition. For replication, the number of parasites per vacuoles were counted. At least 100 vacuoles were counted.

In the case of RHDiCreΔ*ku80* floxed strains, loxP*cgp*-Halo parasites were used for invasion/replication assays, loxP*slf*-Halo were used for invasion assays, and loxP*slf*-mCherry were used for replication assays. Parasites were pre-induced for 96 h ±50 nM rapamycin. Parasites were then mechanically egressed, and 5 × 10^6^ parasites (for invasion assays) or 4 ×10^6^ (for replication assays) were inoculated in each well and left to invade for 1 h.

For invasion assays, parasites were allowed to settle on ice for 10 min, and then allowed to invade for 1 h at 37 °C before fixation. Subsequent IFAs were done following the same protocol as the invasion assays done on the sCas9 parasite strains. A minimum of 150 parasites were counted to calculate the percentage of invaded parasites.

For replication assays, samples were washed 3 times with DMEM to remove non-attached parasites and left at 37 °C for another 24 h. Samples were fixed with 4% PFA and labelled with α-GAP45 (loxP*cgp*-Halo) or IMC1 and α-RFP (loxP*slf*-mCherry).

LoxP*cgp*-Halo parasites were pre-incubated with HaloTag Oregon Green dye (0.2 μM) for 1 h, whereas loxP*slf*-Halo parasites were incubated with Halo Janelia 646 (20 nM) for 15 h prior to the start of the invasion assay.

These experiments were carried out in triplicate and a minimum of 100 parasites/vacuoles were counted (n = >100) In case of rapamycin induced floxed parasites (cKO), only parasites/vacuoles lacking signal for SLF or CGP were included in the counting.

### Plaque assay

A total of 500-1000 parasites per well were inoculated into confluent HFFs in 6 well-plates and incubated for 6 days ± 50 nM rapamycin as previously described ^32^.

In case of GABA (Tocris, 0344) or Gabapentin (Sigma-Aldrich, G154) plaque assays, media was supplemented with different concentrations of GABA (100 mM stock concentration) or Gabapentin (50 mM stock concentration).

Images were taken using the LAS X Navigator software and a Leica DMi8 Widefield microscope using 10× objective (Microsystems). Starting in the middle of the well, an area of 12 x 12 fields was imaged. Focus maps were created and autofocus controls were applied for taking the final images. After acquisition of the images, “mosaic merge” processing tool in LAS X software was used for merging the pictures into one big final image.

### Egress assays

For egress assays depicted in Supplementary Fig. 3, induced (50nM Rapamycin) and non-induced RHsCas9-CbEm-adf/sag1 parasites were grown for 48h. Egress was then induced by incubating parasites with 2μM Ci A23187 for 8min under normal culturing conditions. Subsequently, parasites were fixed with 4%PFA for 20min and α-SAG1 or α-GAP45 were used for parasite visualisation.

1×10^5^ sCas9 background parasites were grown on HFF cells incubated with ± 50 nM rapamycin for 4 h. They were then washed with DMEM three times to remove rapamycin and any extracellular parasites. Parasites were maintained for another 44 h in the incubator at 37 °C before inducing egress.

In case of parasites floxed in the DiCre strain, parasites were pre-treated ± 50 nM rapamycin for 24 h and 3 × 10^5^ were inoculated onto confluent HFF cells, incubated for 1 h at 37 °C, then washed 3 times with PBS and maintained at 37 °C for 32 h before inducing egress. Halo tagged parasites were pre-incubated with HaloTag Oregon Green (0.2 μM) or Janelia Fluor 646 (20 nM) for 1 h and washed three times with PBS before inducing egress. To induce parasite egress, media was exchanged with pre-warmed DMEM without serum but supplemented with various inducers for different lengths of time (2 μM Ci A23187 (Sigma-Aldrich, C7522-1mg) for 5 min, 50 μM BIPPO (a phosphodiesterase inhibitor that stimulates microneme secretion^19^; Thompson Lab) for 5 min, or 125 μM Propranolol hydrochloride (Merck, 40543) for 7 min).

After induction, sCas9 expressing parasites were fixed with 4% PFA and the number of egressed and non-egressed vacuoles counted. DiCre-expressing parasites were fixed with either 4% PFA or 100% methanol. α-SAG1 or α-GAP45 antibodies were used for the visualisation of parasites. In case of rapamycin induced floxed parasites, only vacuoles lacking signal of the respective protein was considered in the counting (cKO). At least 100 vacuoles were counted for each condition and replicate and the percentage of the egressed vacuoles was calculated.

For time-lapse images, floxed parasites expressing CbEm (loxP*cgp*-Halo/CbEm and loxP*slf*-Halo/CbEm) were treated ± 50 nM rapamycin for 24 h and then mechanically released and inoculated onto glass-bottom live cell dishes, and cultured for a minimum of 32 h before inducing egress. Halo-tagged parasites were pre-incubated with Janelia Fluor 646 (20 nM) for around 5 h followed by washing 3 times with PBS and then incubated with normal media for at least 1 h before egress induction. Dishes were placed in the pre-warmed chamber of Leica DMi8 microscope and media was exchanged with complete Fluorobrite DMEM (ThermoFischer Scientific, A1896701) containing the respective inducers. Videos were taken with a 63x oil objective at 0.33 frames per second (FPS). Videos were recorded in triplicate per condition as a minimum. In case of conditional knockouts, only vacuoles lacking the signal for SLF or CGP were recorded (cKO).

For calculating the dynamics of CbEm after stimulating egress chemically, regions of interest (ROI) were drawn around the apicoplast region, the region between basal CbEm labelling and the apicoplast (termed nuclear region), the *T. gondii* cell, and a background region outside the vacuole. Relative intensity of the CbEm in apicoplast region was then determined as:

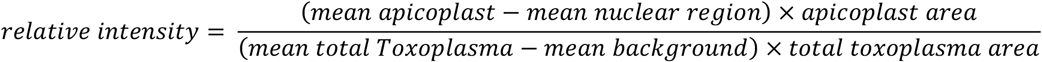

Relative intensity of the CbEm in basal pole region was then determined as:

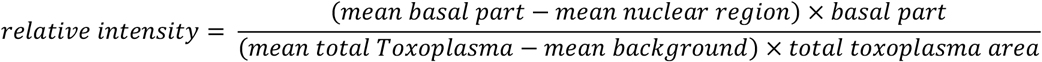

where mean was defined as:

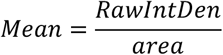

To check PVM integrity, loxP*cgp*-Halo/CbEm and loxP*slf*-Halo/CbEm were treated with ± 50 nM rapamycin for 24 h and transfected with the vector pTub-*sagf*ΔGPI-dsRed. 48 hours later, egress was induced with 50 μM BIPPO and recorded as described above. Over 10 egress events were recorded for each condition.

### Trail deposition assay and live gliding assay

For trail deposition assays, induced (50 nM rapamycin, 72 h for loxP*cgp*-Halo and 96 h for loxP*slf*-mCherry) and non-induced (72 h for loxP*cgp*-Halo and 96 h for loxP*slf*-mCherry) parasites were washed, mechanically egressed via 26-gauge needles and filtered through 3 μm filters. Parasites were then centrifuged at 1000 x g for 5 min at room temperature and the pellet was resuspended in pre-warmed endo buffer (44.7 mM K_2_SO_4_, 10 mM MgSO_4_, 100 mM sucrose, 5 mM glucose, 20 mM Tris, 0.35% w/v BSA, pH 8.2) at a concentration of 2×10^6^ parasites/ml. 1 ml of this mixture was added to a FCS coated glass-bottom live cell dish and incubated for 15 min at room temperature. Endo buffer was gently replaced with 1ml of pre-warmed sterile gliding buffer (1mM EGTA and 100mM HEPES in HBSS solution). Parasites were incubated for 20 min at 37 °C and then fixed with 4% PFA for 20 min. Parasites were stained with *α-Toxoplasma gondii* antibody (Abcam; see Supplementary table 6) without permeabilisation. 15 random fields of view were imaged and the total number of trails parasites left were counted.

For live gliding assays, to measure parasite gliding kinetics, time-lapse videos were taken with a 63× objective at 2 FPS using a Leica DMi8 microscope using DIC. After 20 minutes of recording per condition, a Z-stack image of the fluorescence channel targeting the protein of interest was taken to distinguish the cKO from the non-induced parasites. Only cKO were considered for the analysis for rapamycin induced parasites. Halo tagged parasites were pre-incubated with Janelia Fluro 646 dye (20 nM) or HaloTag TMR (500 nM) at least 2 hour as described above before performing live gliding assays. Parasite motility was analysed by manual tracking plugin tool by Icy software. All the assays were done in Ca^2+-^free gliding buffer unless otherwise indicated.

For trail deposition and gliding assays with 2 μM Ci A23187, compounds were added to the gliding buffer described above.

### Microneme secretion assay

Microneme secretion assay protocol was adapted from Bisio *et al.* (2019)^20^. Parasites treated for 72 h ± 50 nM rapamycin were mechanically released by 26-gauge needles and washed once with cold PBS before being resuspended in pre-warmed intracellular buffer (5 mM NaCl, 142 mM KCl, 1 mM MgCl2, 2 mM EGTA, 5.6 mM glucose, 25 mM HEPES, pH 7.5). Parasites were then incubated ± 200 nM A23187, 5 μM BIPPO or DMSO for 30 min at 37 °C. Afterwards, the supernatant was collected and further centrifuged, followed by Western blot analysis (WB). The parasite pellets were washed once with cold PBS followed by centrifugation before performing Western blot analysis. For WB, 6% stacking and 10% resolving gel were used. Antibodies used to label the membranes are summarised in Supplementary Table 6. Stained membranes were imaged using Odyssey CLX-1849 (LI-COR).

### Imaging processing

All images and movies were processed using Fiji software 2.1.0 and/or Icy Image Processing Software 1.8.6.0. With the exception of images of parasites expressing CbEm and time-lapse videos, all widefield images were deconvolved using Huygens essential v18.04.

### Data analysis

All data were plotted by Excel or Graphpad Prism 8.2.1.

## Data availability

All data and genetic material used for this paper are available from the authors on reasonable request. The source data are provided as a Source Data file.

## Author contributions

W.L. identified egress mutants and performed phenotypic assays for SLF and CGP. J.G. designed and performed the phenotypic screen, identified and tagged candidate genes and assisted with phenotyping. J.F.S. established and characterised the sCas9-system in *T. gondii*, designed the sgRNA library and analysed egress for ADF. M.G. assisted with phenotyping and analysis of parasites. J.P. assisted in experiment design and data analysis. M.M. designed and coordinated the project and experiments, analysed the data, contributed resources and wrote the paper. E.J-R. designed and coordinated the project and experiments, analysed the data, and wrote the paper.

## Acknowledgement

We thank all colleagues, who contributed antibodies and reagents for this study. In particular, we thank the Lourido lab (Whitehead Institute for Biomedical Research) in assisting with the design of the sgRNA library and many useful discussions. W.L. is funded via a CSC fellowship (201806910075). This project is funded within the DFG Priority Programme SPP2225, Project ME 2675/7-1.

## Competing interests

The authors declare no competing interests.

## Notes

### Competing Interest Statement

The authors have declared no competing interest.

