## Supplementary figures for "A phenotypic screen using splitCas9 identifies essential genes required for actin regulation during host cell egress and invasion by *Toxoplasma gondii*"

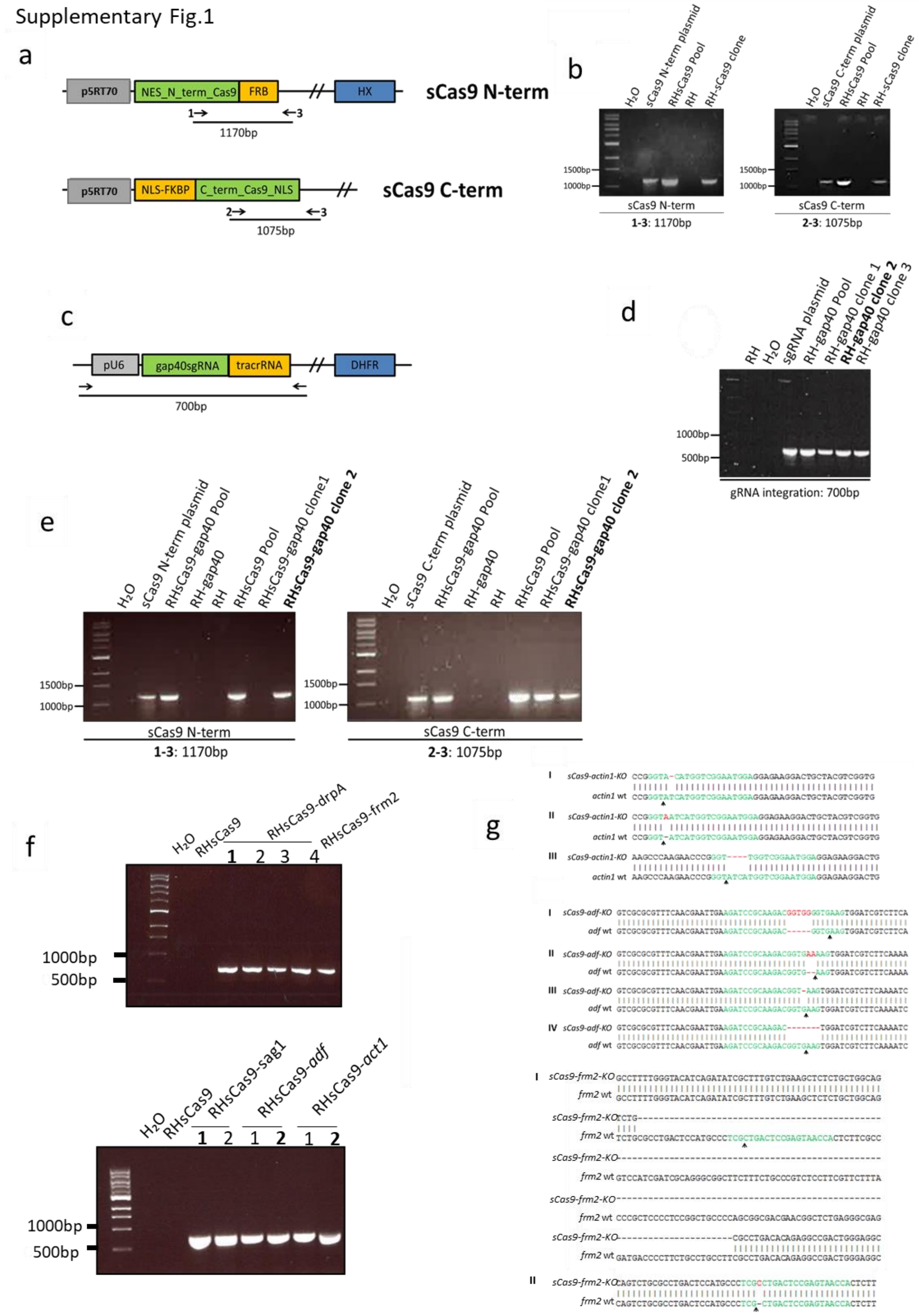


**Supplementary Fig.1 Generation of parasite strains RHsCas9, RH-gap40, RHsCas9-gap40, RHsCas9-sag1, RHsCas9-drpA, RHsCas9-act-1 and RHsCas9-adf. a**, Scheme of expression cassettes for the N- and C-termini of the Cas9 enzyme (split 4 variant, see [4]). Arrows indicate PCR amplicon for verification of plasmid integration (see (b)). **b**, Analytical PCR confirming integration of sCas9 plasmids into the genome of indicated parasites. **c**, Scheme of the expression cassette for the single-guide RNA (sgRNA, here for targeting of gap40). Arrows indicate PCR amplicon for verification of plasmid integration. **d**, Analytical PCR confirming integration of gap40-sgRNA-plasmid into the parasite genome. **e**, Analytical PCRs confirming integration of sCas9 plasmids into the RH parasites. **f**, Analytical PCRs confirming integration of indicated sgRNA-plasmids into the parasite genome. **g,** Validation of specific introduction of indels at the sgRNA cut side in indicated parasites. Cultures were induced with 50 nM rapamycin for 1 h. Parasites were grown for 48 h prior to gDNA collection. The sgRNA cut site was amplified by PCR and sequenced. Red letters represent nucleotide insertion in the mutant strain, causing a frame shift and, thus, the functional disruption of the indicated gene. Black arrows indicate the predicted cut site.

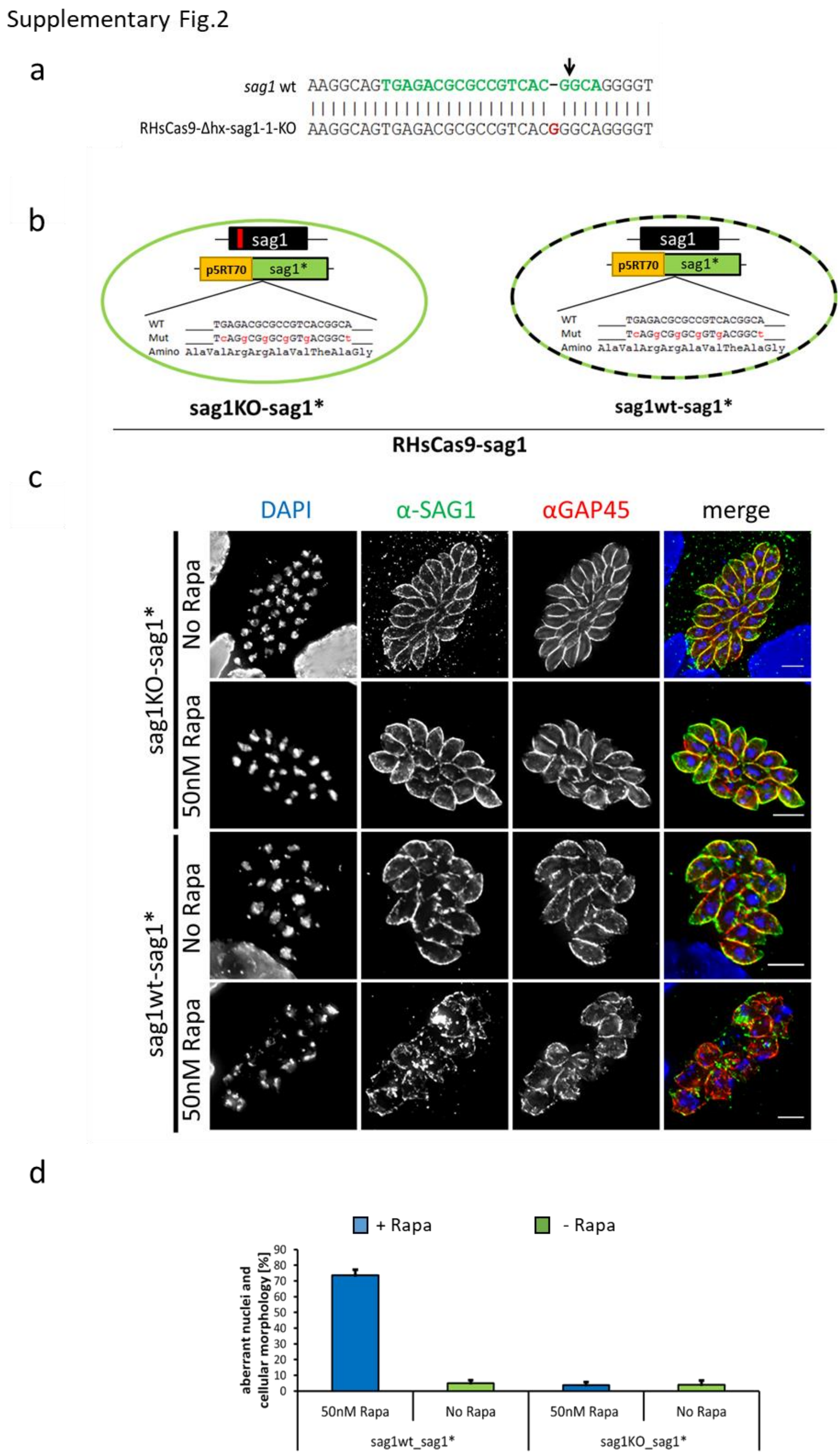


**Supplementary Fig.2 The nuclear phenotype is caused by Cas9-mediated double strand breaks.** **a,** Genome sequencing verified disruption of *sag1* in a clonal RHsCas9-Δhx-sag1-KO mutant. Green letters indicate sgRNA sequence. Red letters represent nucleotide insertions upon disruption of *sag1*, causing frame shifts. The black arrow indicates the predicted cut side. **b,** Schematic of the strains sag1KO-sag1* and sag1wt-sag1*. Both strains were generated in the RHsCas9-sag1 background. The endogenous sag1 gene in the sag1KO-sag1* line has been disrupted by prior sCas9 activation. The endogenous sag1 gene of sag1wt-sag1* is intact. The additional sag1 copy has been modified as indicated (red letters) to be resistant to sag1sgRNA recognition. **c,** IFA depicting the impact of sCas9 activation on sag1KO-sag1* and sag1wt-sag1* parasites. Parasites were induced with 50 nM rapamycin for 1 h or in the absence of rapamycin. Parasites were grown for 48 h prior to IFA analysis using indicated antibodies. Scale bars are 5µm. **d,** Quantification of vacuoles displaying aberrant nuclei and cellular morphology after sCas9 activation at 48h post inoculation. For each condition at least 100 vacuoles were counted (total n⩾300).

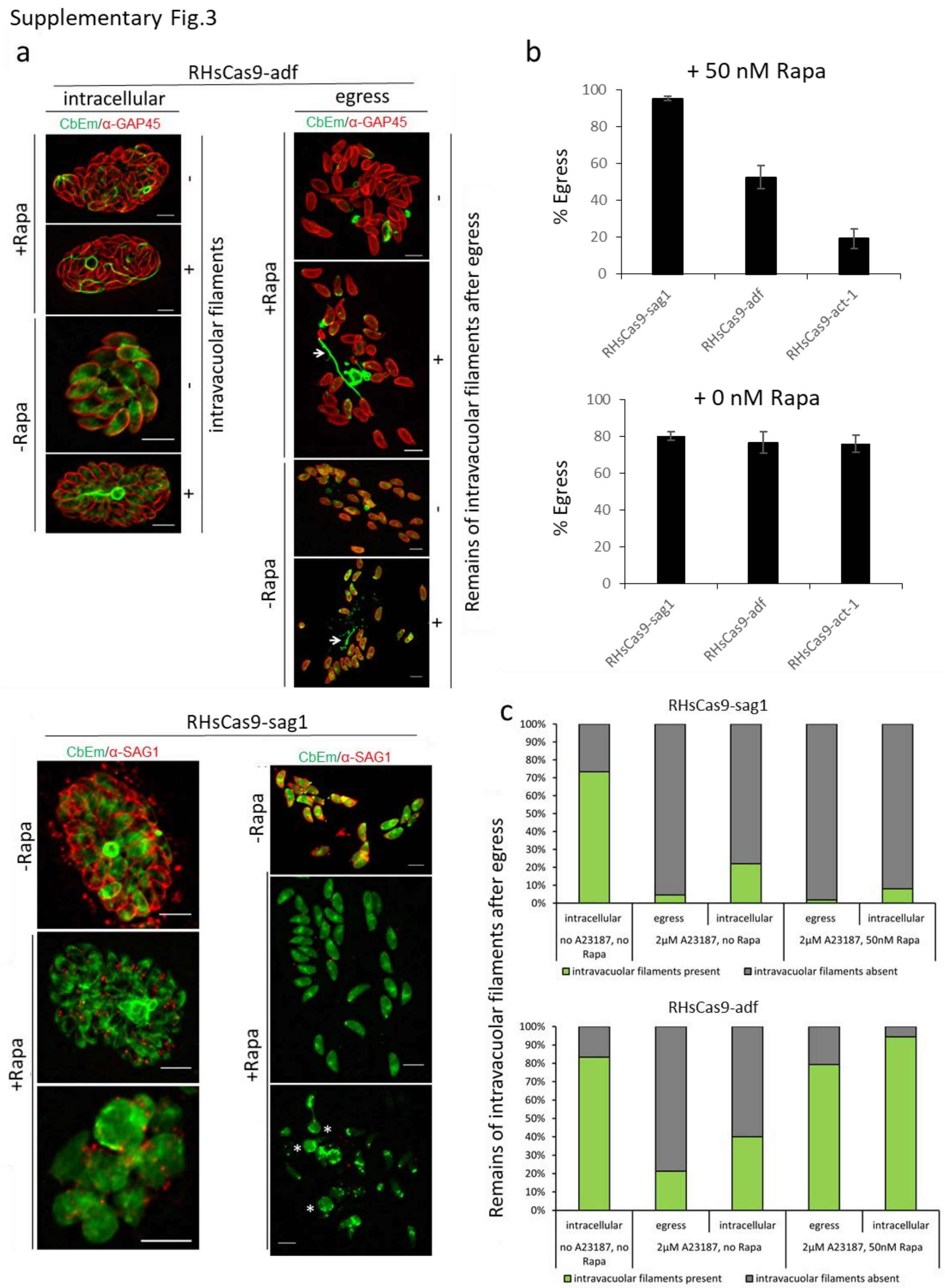


**Supplementary Fig.3 Characterisation of egress in parasites with depleted ADF. a,** Images depict intracellular and egressed vacuoles with or without intravacuolar filamentous actin structures. For this experiment, indicated parasite were induced (50 nM rapamycin, 1h) or non-induced and grown for 48h. Egress was then induced by incubating parasites with 2 µM calcium ionophore (Ci) A23187 for 8 min. After fixation with 4% PFA, parasites were stained for GAP45 by IFA. Scale bars are 5 µm. **b,** Rapamycin induced (50nM, 1h, upper panel) and non-induced parasites (lower panel) were grown for 48 h. Egress was then triggered by incubating parasites with 2 µM Ci A23187 for 8 min. Rate of egress of indicated induced parasites (upper panel) was normalised to egress of the corresponding non-induced parasite population (lower panel). For each condition 100 vacuoles were counted (total n=300) In one of the three biological repeats for RHsCas9-adf, parasites were induced for 48h and egress was initiated with Ci A23187 for 5mins. **c,** Analysis of the abundance of large intravacuolar filamentous actin structures after induction of egress in sag1-wt/KO and adf-wt/KO parasites. See (a) for representative images. Analysis is based on three independent experiments. For RHsCas9- *sag1*: no A23187/no Rapa: n=300; 2 µM A23187/no Rapa: n=241 (egress), n= 59 (intracellular); 2 µM A23187/50 nM Rapa: n=225 (egress), n=75 (intracellular). For RHsCas9- *adf*: no A23187/no Rapa: n=300; 2 µM A23187/no Rapa: n= 230 (egress), n= 70 (intracellular); 2 µM A23187/50 nM Rapa: n=121 (egress), n= 179 (intracellular).

Supplementary Fig.4

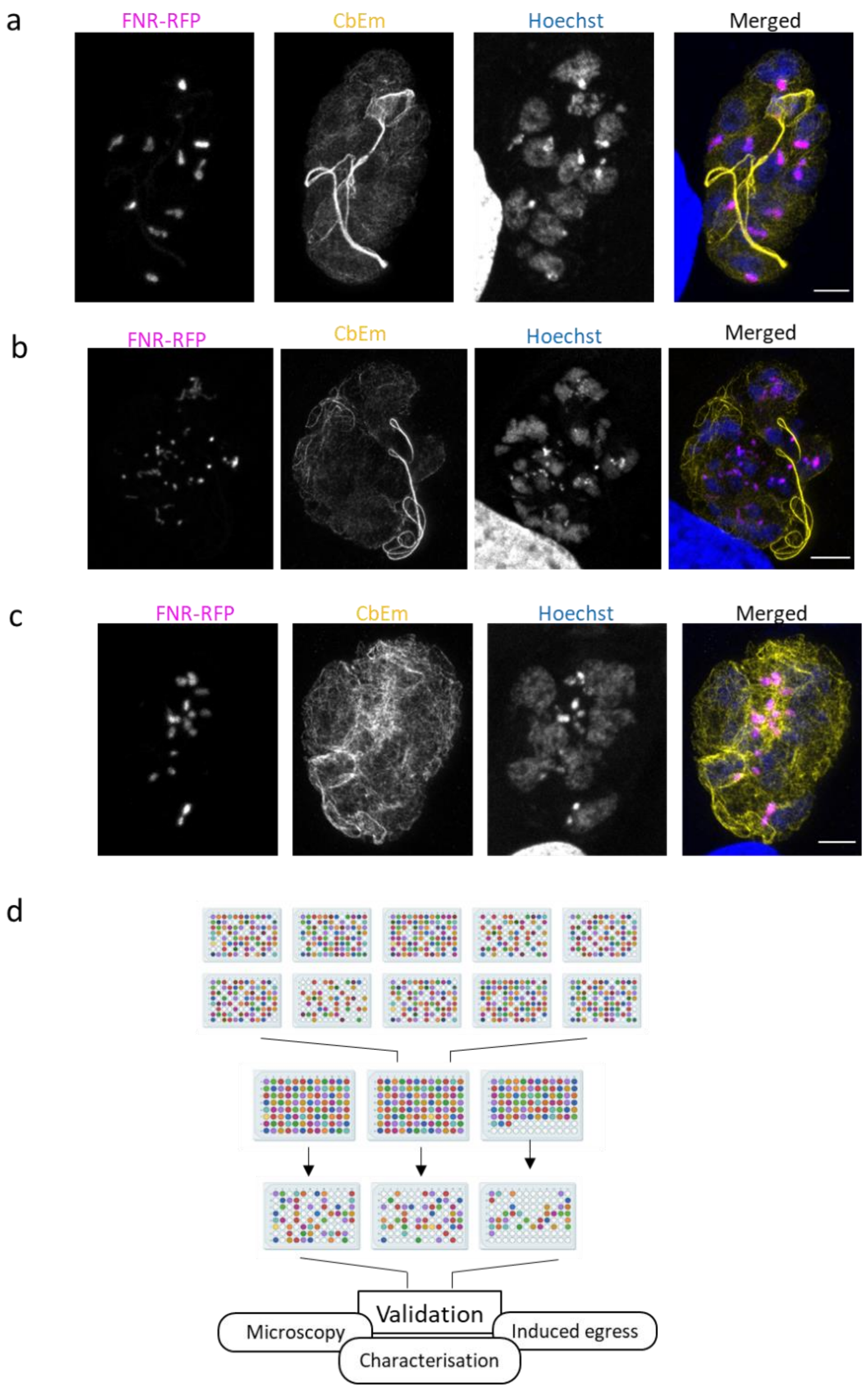


**Supplementary Fig.4 Selection of candidates and super-aberrant phenotype of parasites with multiple sgRNAs. a,** Parasites containing the sgRNA targeting *prf* (positive control for F-actin dynamic phenotype) were isolated multiple times and presented a thick, extensive intravacuolar network. Scale bar is 3 µm. This phenotype is very similar to the one observed after the depletion of ADF. **b,** Parasites containing the sgRNA targeting *gap40* (positive control for replication phenotype). The identified phenotype is identical to the one reported previously [7]. Scale bar is 5 µm. **c,** Some of the most impressive phenotypes discovered were caused by multiple integration of sgRNA targeting different genes. Super-resolution image of a representative clone is shown. Scale bar is 3 µm. These clones were omitted from further analysis. **d,** Schematic for the selection of candidate genes.

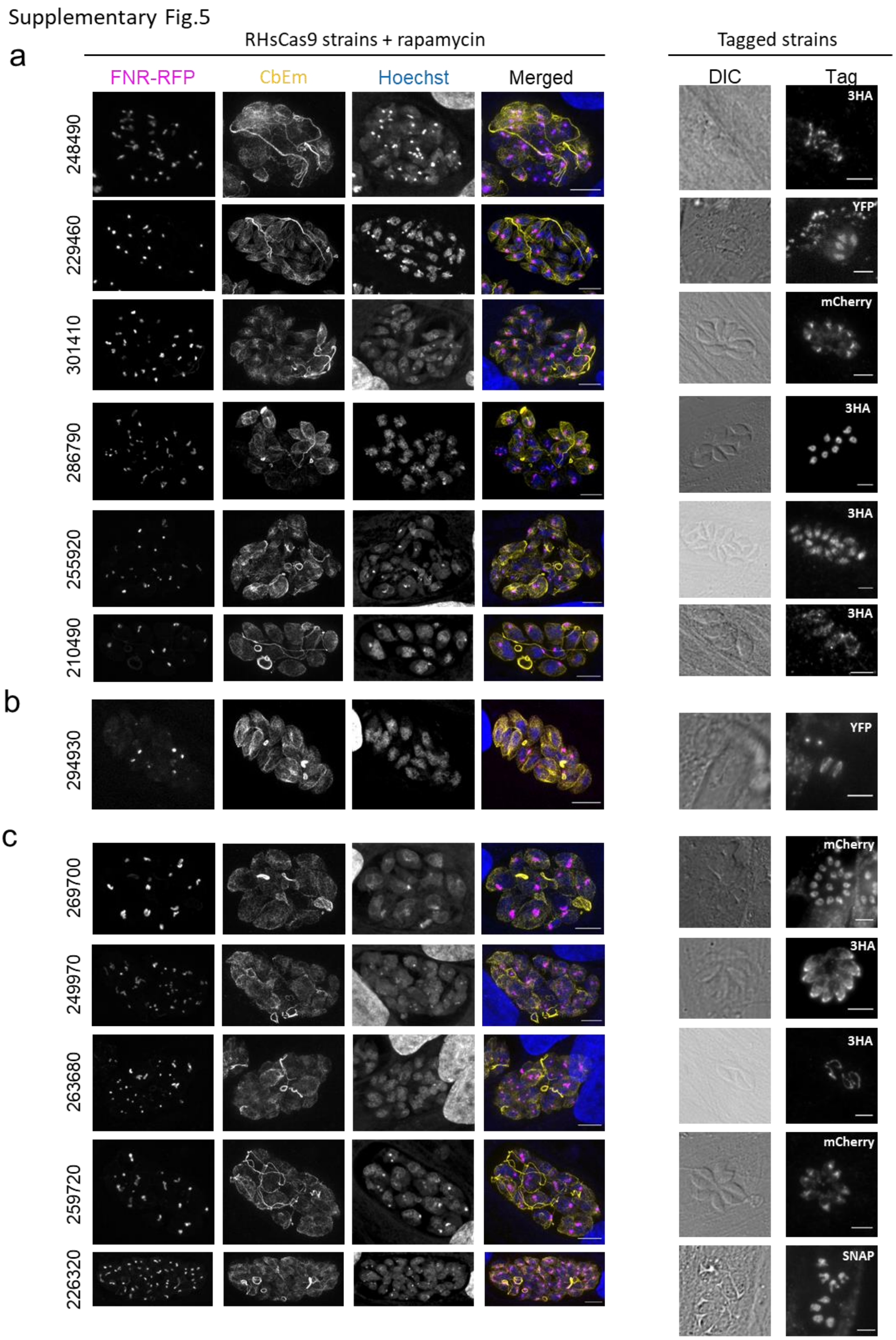


**Supplementary Fig.5 Mutants with (slight) changes in F-actin localisation and apicoplast maintenance. a,** Mutants with F-actin phenotype. **b,** mutants with apicoplast phenotypes. **c,** mutants with both F-actin and apicoplast phenotypes. Left images: Images of sCas9-CbEm-FNR clones containing sgRNAs targeting indicated genes. Parasites were induced for 48 h with rapamycin before fixing and imaging. CbEm, STED images. FNR and Hoechst are confocal images. Right images: Widefield images of RHΔ*ku80* parasites with the respective candidate gene endogenously tagged as indicated. Scale bars are 5 µm.

Supplementary Fig.6

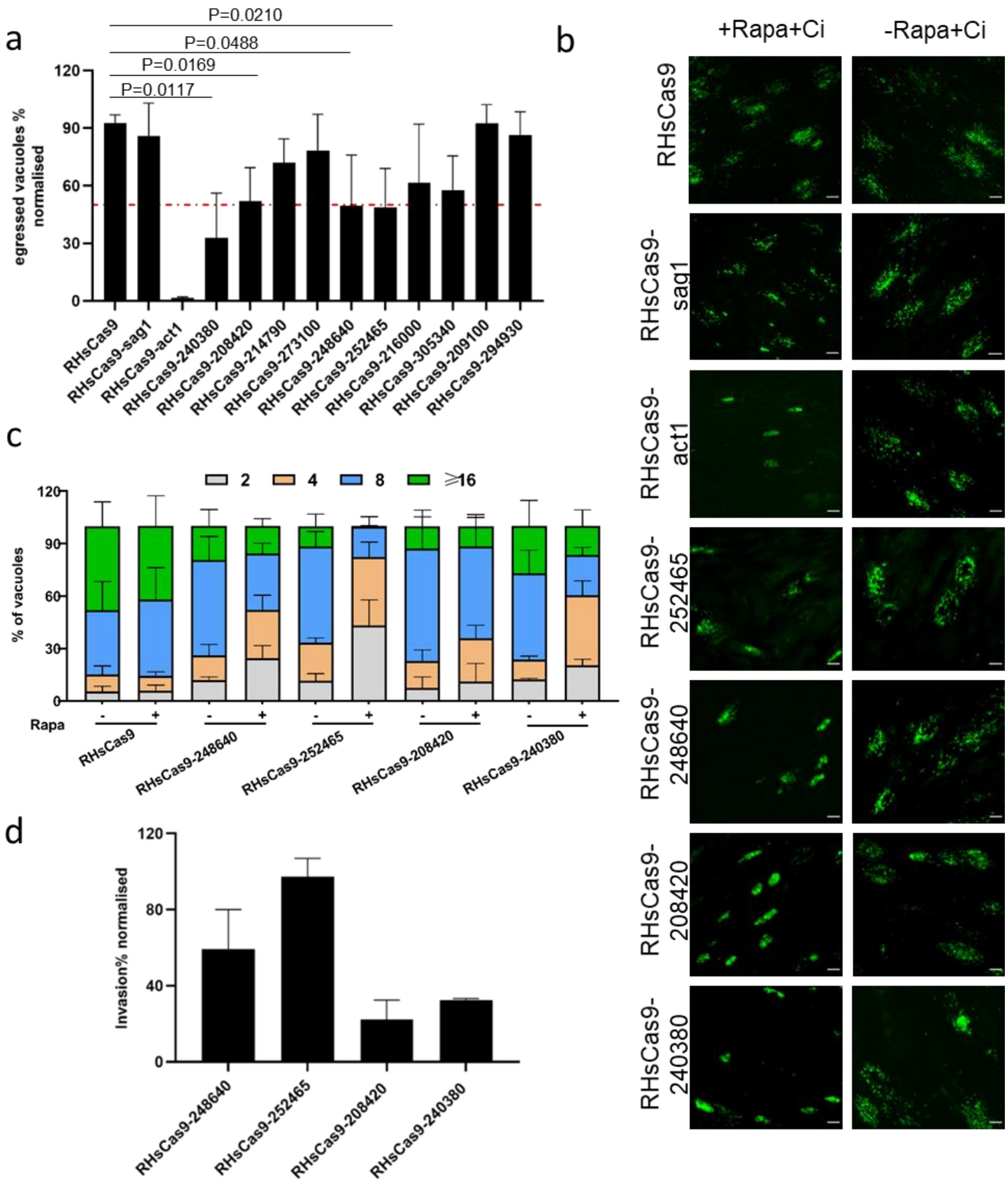


**Supplementary Fig.6 Validation of candidates. a,** Quantification of induced egress in RHsCas9-CbEm-FNR-RFP isolated clones with 2 µM of calcium ionophore (Ci) A23187. Results of the rapamycin (R) induced parasite (+R+Ci) were standardised to the non-rapamycin treated condition (-R+Ci). Red dash line marks reduction of egress to 50% when compared to non-induced parasites. 4 candidates were selected after this analysis. Unpaired two-tailed Student's *t*-test were calculated and resultant P value is shown in the graph. For each condition, at least 100 vacuoles were counted (total n≥300) **b,** Representative images of the 4 candidate genes and control parasites, as quantified in (a). Scale bar, 30 µm. **c,** 24-hour replication assay. Parasites were inoculated onto HFF, washed after an hour and incubated for 24 h. Vacuoles were counted and number of parasites per vacuole was determined. Quantification of parasite replication showed a delay in replication for TGGT1_252465, which was omitted from further analysis. For each condition, at least 100 vacuoles were counted (total n≥300). **d,** Quantification of invasion of indicated parasites was standardised to the non-rapamycin treated condition and normalised to RHsCas9 strain. These assays were performed in triplicates. Standard deviation error bars are shown in their respective graph. For each condition, at least 100 vacuoles were counted (total n≥300).

Supplementary Fig.7

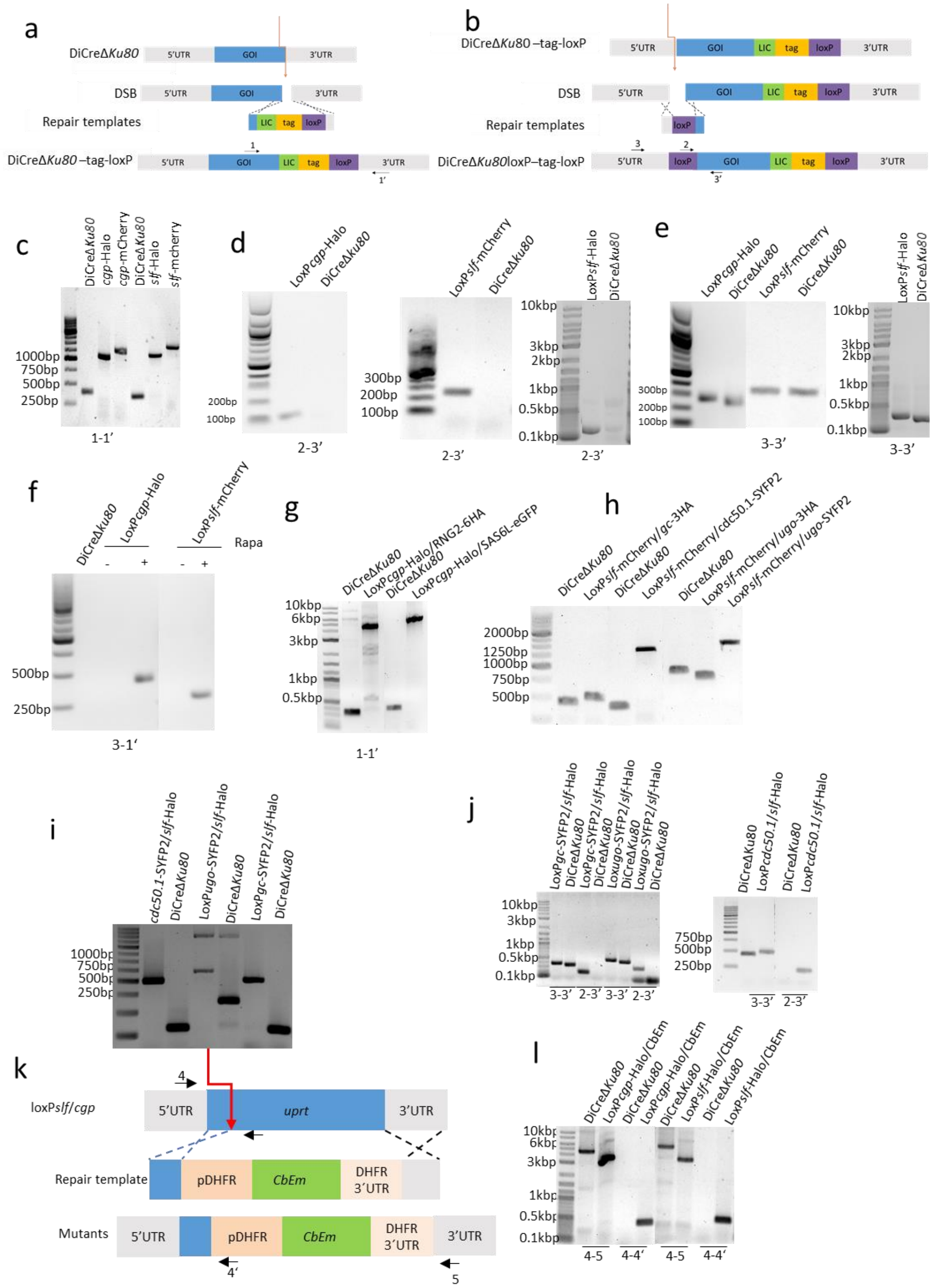


**Supplementary Fig.7 Generation of parasite lines in DiCre background parasites. a,** C-terminal tagging scheme. Donor DNA containing respective tag, loxP sequence and 50 nucleotides of homologous flanking sequence was co-transfected with gRNA-Cas9YFP vector. Primers 1 and 1' bind outside of the homologous region incorporated in donor DNA and were used to verify insertion of the tag. Orange arrow indicates the targeted region of the sgRNA. **b,** Generation of floxed lines. Insertion of upstream loxP sequence before the open reading frame at 5'UTR region was achieved by co-transfection of gRNA-Cas9YFP together with a long oligo containing a loxP sequence flanked by 50 nt of homology to the targeted region. Primers 2 and 3' were used for verification of loxP integration. Primers 3 and 3' bind outside of the homologous region flanking the loxP and were used for sequencing of the insertion. Primers 3 and 1' were used for confirmation of gene excision. Orange arrow indicates the targeted region of the sgRNA. **c,** Analytical PCR shows the correct integration of tags in the indicated parasite lines. **d,** Integration PCR indicates the loxP insertion at the 5'UTR in the indicated parasite lines. **e,** Genotyping PCR verifying the loxP insertion at 5'UTR in the indicated parasite lines. These amplicons were sent for sequencing to confirm correct sequences. **f,** PCR indicating excision of the floxed gene. **g,** Correct integration of 6HA and eGFP for RNG2 and SAS6L in loxP*cgp*-Halo parasite lines, respectively. **h,** PCR shows the correct integration of tags in the floxed parasite lines. **i,** Genotyping PCR indicates successful tagging of *gc*, *cdc50.1*, and *ugo*. **j,** Genotyping PCR indicates the loxP insertion at 5'UTR in the indicated parasite lines**. k,** Scheme for replacing UPRT locus with CbEm driven by Tg*dhfr* promotor. Orange arrow indicates the targeted region of the sgRNA. Primers 4 and 4' were used to verify insertion of the tag. Primers 4 and 5 were also used to verify insertion of the tag. **l,** PCRs showing integration and for genotyping of CbEm in floxed parasite lines.

Supplementary Fig.8

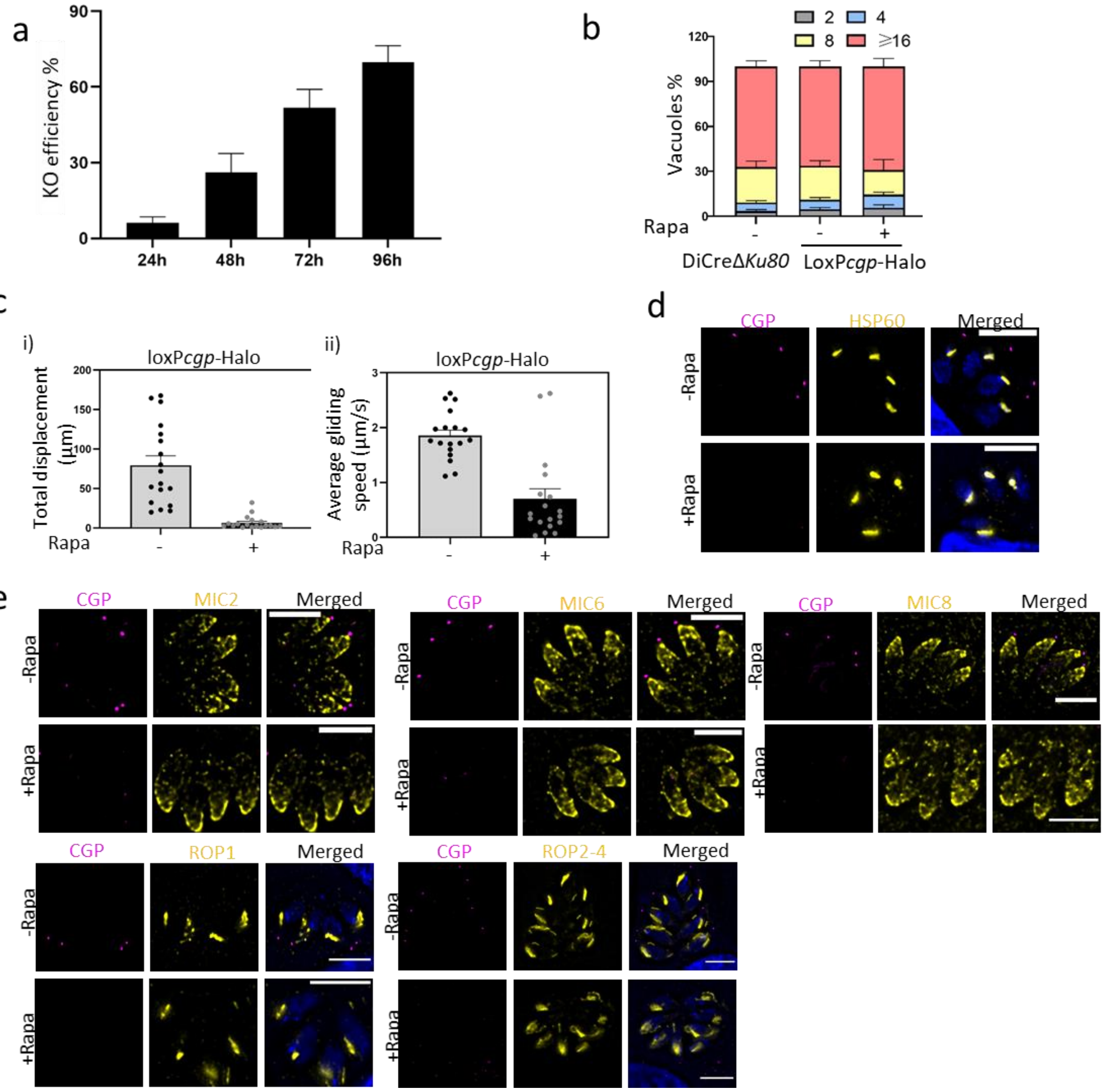


**Supplementary Fig.8 Loss of CGP doesn't affect parasite morphology or secretory organelles. a,** Quantification of the percentage of vacuoles with loss of CGP signal. loxP*cgp*-Halo parasites were treated with 50 nM rapamycin for 24 h, 48 h, 72 h, and 96 h**.** At least 100 vacuoles were counted (total n≥300). **b,** Parasites lacking *cgp* showed a normal intracellular replication. For each condition, at least 100 vacuoles were counted (total n≥300). **c,** The gliding length (i) and average gliding speed (ii) of parasites capable of gliding (helical or circular movement) were measured by 18 tracked parasites. Motility was analysed by manual tracking using the plugin by Icy software. Data were presented Mean ± SEM. **d-f)** Localisation of apicoplast, microneme and rhoptry proteins is not affected upon deletion of *cgp*. Parasites were pre-treated ± 50 nM rapamycin for 1 hour and imaged 72 hours later. HSP60: marker for apicoplast. MIC2, MIC6, and MIC8: marker for microneme proteins. ROP1 and ROP2-4: marker for rhoptry proteins. Scale bar, 5 µm.

Supplementary Fig.9

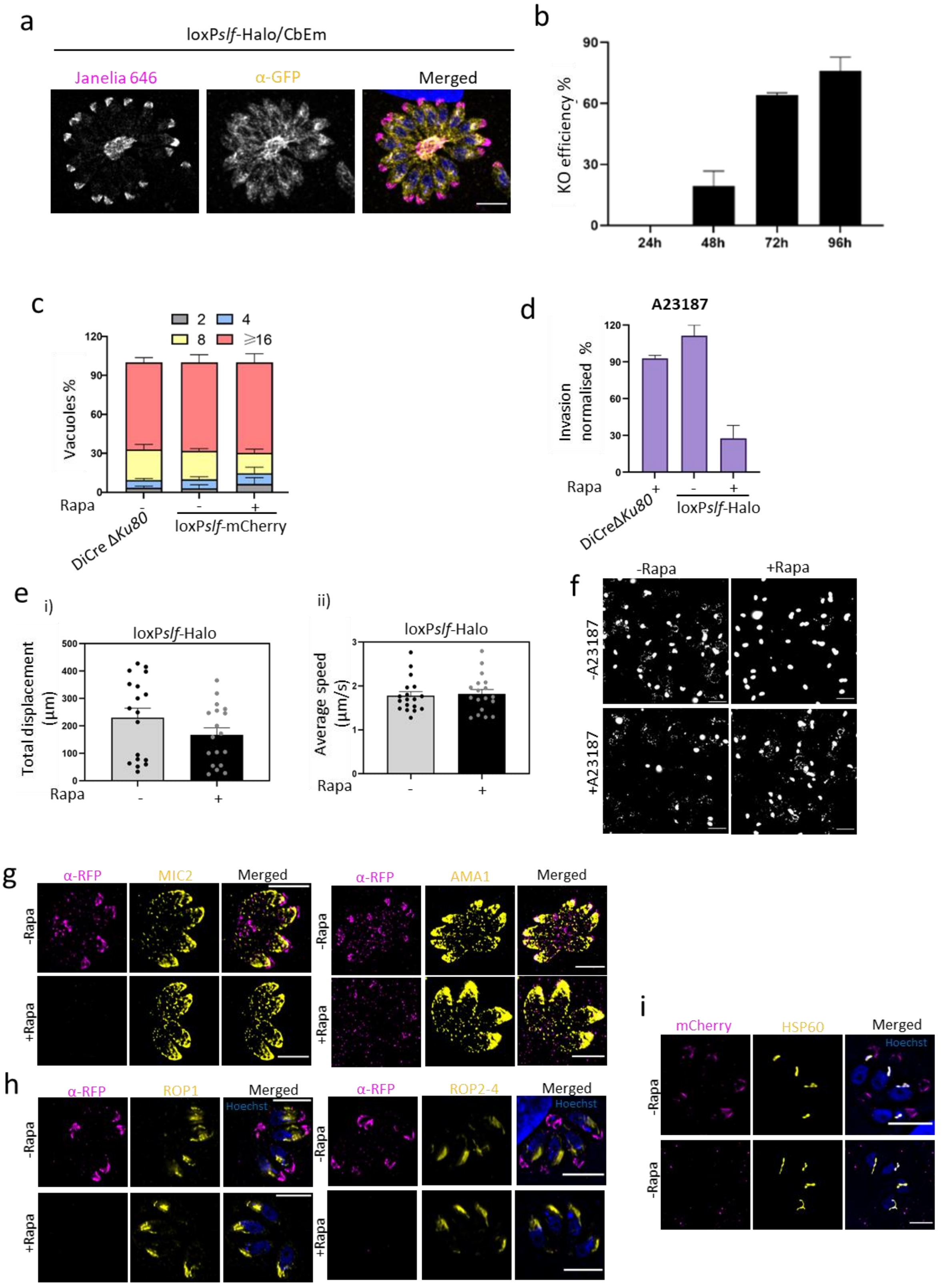


**Supplementary Fig.9 Deletion of SLF does not interfere with microneme or rhoptry trafficking. a,** SLF localised at the apical tip and the intravacuolar network, as shown by co-localisation with CbEm (labels F-actin). Scale bar, 5 µm**. b,** Quantification of the percentage of vacuoles without SLF signal in loxP*slf*-mCherry parasites treated with 50 nM rapamycin for 24 h, 48 h, 72 h, and 96 h. At least 100 vacuoles were counted per replica (total n≥300). **c,** Parasites lacking *slf* showed a normal intracellular replication. For each condition, at least 100 vacuoles were counted (total n≥300). **d,** Invasion attachment assay in the presence of 2 µM calcium ionophore A23187 after 96 hours post induction. Data was normalised to DiCreΔ*Ku80* parasites and presented as mean ± SD. 3 biological replicates were done. For each condition, at least 150 vacuoles were counted (total n≥450). **e,** The gliding length (i) and average gliding speed (ii) of parasites capable of gliding (helical or circular movement) was measured by 18 tracked parasites in the presence of 2 µM Ci A23187. Movements was analysed by manual tracking using the Icy software plugin. Data presented is mean ± SEM. **f.** Trail deposition assay of loxP*slf*-mcherry parasites pretreated ± 50 nM rapamycin for 96 hours stimulated with or without 2 µM Ci A23187 indicates that the number of cKO parasites capable of initiating motility was increased when induced with Ci A23187. Therefore reduction in overall gliding is likely caused by a block in microneme secretion (see Fig.3g). **g-i,** Apicoplast, microneme and rhoptry proteins are not affected upon deletion of *slf*. Parasites were pretreated ± 50 nM rapamycin for 1 hour. HSP60, marker for apicoplast. MIC2 and AMA1: marker for microneme proteins. ROP1 and ROP2-4: marker for rhoptry proteins. Scale bar: 5 µm.

Supplementary Fig.10

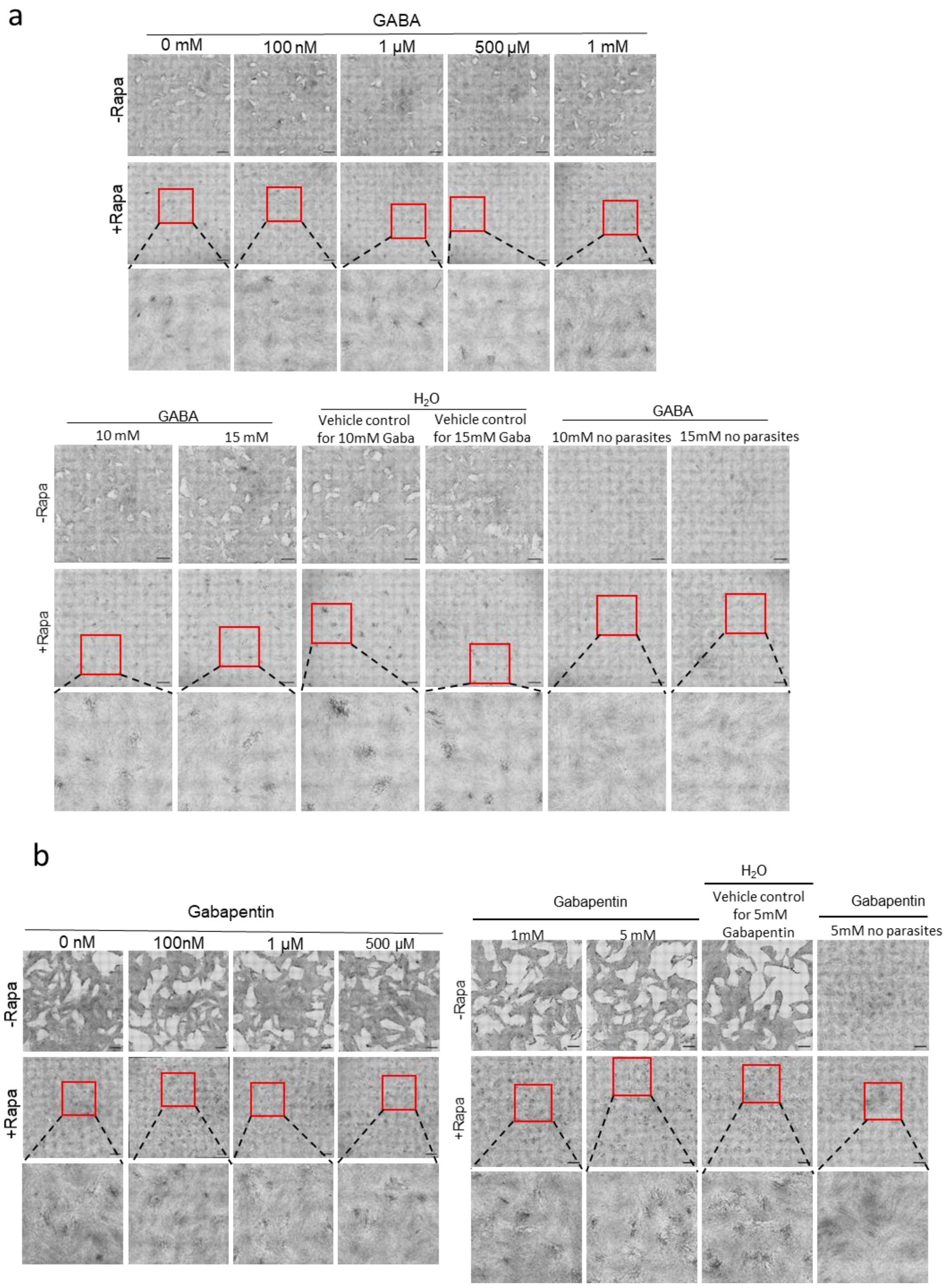


**Supplementary Fig.10. GABA and GABA analogues effect on parasites with depleted SLF**. Plaque assay of loxP*slf*-mCherry supplemented with different concentration of GABA (a) and Gabapentin (b) indicating SLF does not play a role in GABA signalling in *Toxoplasma*. Scale bar: 1.5 mm.

### Supplementary movies:

**Movie_S1. Time-lapse video microscopy of gliding parasites. a,** Non-induced loxP*cgp*-Halo parasites gliding normally. **b-c**, Gliding in pre-induced loxP*cgp*-Halo parasites with 50nM rapamycin (only parasites not expressing CGP were analysed). **b,** Non-gliding *cgp*-cKO. **c,** Gliding *cgp*-cKO. Note movement was slow and the distance covered was minimal. **d,** Non-induced loxP*slf*-Halo parasites gliding normally. **e-f,** Gliding in induced loxP*slf*-Halo parasites with 50nM rapamycin (only parasites not expressing SLF were analysed). **e,** In most cases, *slf* cKO parasites seemed to float over the surface. **f,** Some *slf* cKO parasites attached on the FCS-coated surface. However, no moment was observed. Time indicated in minutes : seconds. Scale bars, 5 μm.

**Movie_S2. Time-lapse video microscopy of loxP*slf* gliding parasites in the presence of 2 μM Ci A23187. a,** Non-induced with rapamycin, loxP*slf*-Halo parasites glided normally. **b,** Induced KO parasites (only parasites not expressing SLF were analysed; *slf*-cKO) can glide normally upon addition of Ci A23187. Time indicated in minutes : seconds. Scale bars: 5 μm.

**Movie_S3. Egress induction in LoxP*cgp*-Halo expressing CbEm parasites (loxP*cgp*-Halo/CbEm. a-c**, Non-rapamycin induced parasites. **a,** normal egress after addition of BIPPO. **b,** parasites egress normally after addition of calcium ionophore (Ci) A23187. **c**, egress after addition of propranolol. **d-f**, Parasites induced with rapamycin prior to the egress induction with different compounds (only parasites not expressing CGP were analysed). **d,** although parasites are able to disassemble the F-actin network and the polymerisation centre at the Golgi area disappears, they cannot initiate movement to egress from the host cell after addition of BIPPO. **e,** addition of Ci A23187 does not rescue the egress phenotype observed previously. **f,** upon addition of propranolol hydrochloride, parasites remain inside the host cell. Videos were recorded at 0.33 frames per second. Time indicated in minutes : seconds. Scale bar, 5 μm.

**Movie_S4. Egress induction in LoxP*slf*-Halo expressing CbEm parasites (loxP*slf*-Halo/CbEm. a-c**, Non-rapamycin induced parasites. **a,** normal egress after addition of BIPPO. **b,** parasites egress normally after addition of calcium ionophore (Ci) A23187. **c**, egress after addition of propranolol. **d-f**, Parasites induced with rapamycin prior to the egress induction with different compounds (only parasites not expressing SLF were analysed). **d,** parasites are unable to initiate egress after addition of BIPPO. Note that the intravacuolar network and the polymerisation centre at the Golgi remains intact. **e-f,** addition of Ci A23187 rescue partially the egress phenotype observed previously. Note parasites are able to disassemble F-actin filaments (**e**) but in some cases unable to leave the host cell (**f**). **g,** upon addition of propranolol hydrochloride, parasites remain inside the host cell despite being able to disassemble the filamentous network. Videos were recorded at 0.33 frames per second. Time indicated in minutes : seconds. Scale bar, 5 μm.

**Movie_S5. Parasitophorous vacuole membrane (PVM) integrity of parasites transiently expressing SAG1ΔGPI-dsRed after induction with BIPPO. a-b,** dsRed signal diffuse rapidly after initiation of egress in wildtype parasites (non-rapamycin induced) indicating rupture of the PVM prior egress. **c**, in parasites lacking CGP, PVM lyses but parasites are unable to initiate movement. **d,** depleted SLF prevent lysis of PVM and movement initiation in BIPPO induced parasites. F-actin: yellow. SAG1ΔGPI-dsRed: pink. Time interval between each frame is 2 seconds for non-induced parasites, 5 s for *cpg*-cKO parasites and 10 s for *slf*-cKO. Time indicated in minutes : seconds. Scale bar, 5 μm
